# A Subset of G Protein-Coupled Serotonin Receptor Genes is Linked to a Neuronal Gene Expression Signature and Clinically Favorable Biology in IDH-Mutant Gliomas

**DOI:** 10.64898/2026.08.23.746569

**Authors:** Flávio Leitão Carvalho-Filho, Henrique Ritter Dal-Pizzol, Gustavo R. Isolan, Rafael Roesler

## Abstract

Increasing evidence indicates that neurotransmitter signaling and neuronal interactions are important determinants of glioma biology. However, the clinical and biological significance of serotonin (5-hydroxytryptamine; 5-HT) receptor expression in lower-grade glioma (LGG) remains poorly understood. Here, we investigated G protein-coupled 5-HT receptor genes in LGG using transcriptomic and clinical data from The Cancer Genome Atlas (TCGA-LGG) and Chinese Glioma Genome Atlas (CGGA) cohorts. Initial survival screening identified *HTR1A*, *HTR2A*, *HTR2C*, and *HTR6* as the genes most consistently associated with longer overall survival (OS). Multivariable Cox regression further identified *HTR2A* and *HTR6* as independently associated with longer OS after adjustment for age, sex, tumor grade, and IDH/1p19q molecular subtype. Expression of the four genes was preferentially associated with molecular features of less aggressive gliomas, particularly IDH-mutant tumors. Single-cell RNA-sequencing (scRNA-seq) data supported malignant glioma cells as a major source of their expression, while cell-type deconvolution revealed strong positive associations with neuronal enrichment and inverse associations with stromal and immune signatures. Transcriptome-wide co-expression and Gene Ontology analyses showed that all four receptor genes were associated with neuronal and synaptic programs involving neurotransmitter release, synaptic vesicle function, ion channels, and synaptic signaling. These transcriptional programs were particularly coherent in IDH-mutant gliomas and more heterogeneous in IDH-wildtype tumors. Together, these findings identify a subset of 5-HT receptor genes associated with favorable clinical and molecular features in LGG and suggest that their expression may mark a neuronal/synaptic differentiation state, particularly within IDH-mutant gliomas.

## Introduction

Increasing evidence indicates that brain tumors such as gliomas are not merely passive occupants of the brain but active participants in neural circuits. Bidirectional interactions between glioma cells and surrounding neurons importantly influence tumor initiation, growth, invasion, and therapeutic resistance. Functional neuron-to-glioma synapses are established and become neurochemically and electrically integrated into neural networks, enabling neuronal activity to stimulate tumor progression. These interactions are mediated by multiple neurotransmitter systems and activity-dependent signaling mechanisms, including glutamatergic neurotransmission through α-amino-3-hydroxy-5-methyl-4-isoxazolepropionic acid receptors (AMPARs) and *N*-methyl-D-aspartate (NMDA) receptors, release of brain-derived neurotrophic factor (BDNF), neuroligin-3 and other neuronal growth factors, and gap junction-mediated communication that amplifies signaling within tumor cells (Barron et al. 2025; Monje 2025; Drexler et al. 2025a; Taylor et al. 2023; Venkataramani et al. 2019; Venkatesh et al. 2015; 2019). These discoveries have fundamentally changed our understanding of glioma biology, revealing that neural activity and neurotransmitter receptor signaling constitute major regulators of the malignant phenotype.

The current knowledge of neuron-glioma communication derives predominantly from studies in glioblastoma (GBM), which is the grade IV glioma and the most aggressive form of primary brain cancer in adults. In contrast, comparatively little is known about the relationship between neurotransmitter receptor signaling and tumor biology in lower-grade gliomas (LGGs). These tumors differ from GBM in their molecular landscape, being largely characterized by IDH mutations and distinct developmental trajectories. LGGs typically affect younger adults and follow a slower clinical course, yet almost invariably undergo malignant progression over time (Bready and Placantonakis, 2019; Carriere et al. 2026; Chang et al. 2016; Kihlstedt et al. 2025). Elucidating neurotransmitter-associated transcriptional programs in LGGs may therefore reveal clinically relevant biological mechanisms that differ from those in GBM.

Serotonin (5-hydroxytryptamine, 5-HT) is a major neurotransmitter that regulates multiple aspects of central nervous system development and function, including neuronal excitability, synaptic transmission, synaptic plasticity, learning, memory, and adult neurogenesis (Huang and Kandel; 2007; Ogelman et al. 2024; Udoh et al. 2024; Upreti et al. 2019). Its diverse physiological effects are mediated by a family of fourteen receptor subtypes grouped into seven classes (5-HT1–5-HT7). Most 5-HT receptors are G protein-coupled receptors (GPCRs), with the exception of the ionotropic 5-HT3 receptor (Bockaert et al. 2009; Hoyer and Martin, 1997).

Beyond their established roles in neurotransmission, 5-HT receptors have also emerged as regulators of cell proliferation, migration, differentiation, survival, and angiogenesis in both normal and pathological tissues (Tan et al 2025; Wouters et al. 2009; Zamani and Qu 2012). Accordingly, dysregulation of serotonergic signaling has been implicated in the development and progression of multiple human malignancies, although the specific contribution of individual receptor subtypes appears to be highly context-dependent and remains incompletely understood (Gwynne et al. 2021; Ji et al. 2026; Kolan et al. 2019; Li et al. 2026).

Despite growing evidence linking serotonergic signaling to cancer, comparatively little is known about the transcriptional landscape and clinical significance of 5-HT receptor genes in LGGs. In the present study, we performed a transcriptome-wide analysis of genes encoding a subset of G protein-coupled 5-HT receptors more strongly associated with patient survival in LGG.

## Methods

### Datasets and Gene Expression

The general methods were similar to those described in our previous studies on other synaptic genes (Gaia et al., 2026; Rodrigues et al. 2026) and a workflow of the present study is summarized in Fig. 1. Transcriptomic and clinical data from patients with LGG were obtained from two independent publicly available cohorts: The Cancer Genome Atlas Lower-Grade Glioma project (TCGA-LGG; https://www.cancer.gov/tcga) (Cancer Genome Atlas Research Network; Brat et al. 2015) and the Chinese Glioma Genome Atlas (CGGA; http://www.cgga.org.cn) (Zhao et al. 2021). Both cohorts comprise diffuse gliomas classified as World Health Organization (WHO) grades 2 and 3. Although grade 3 gliomas have historically been grouped with “high-grade” tumors and, in some clinical settings, considered together with glioblastoma (GBM), the TCGA-LGG and CGGA datasets analyzed here were specifically established to encompass diffuse grade 2 and grade 3 gliomas. We therefore adopted the term “lower-grade glioma” (LGG), in accordance with the nomenclature of these datasets and its widespread use in previous transcriptomic studies. Survival analyses included 490 tumors from TCGA and 407 tumors from CGGA. Gene expression was analyzed across molecularly defined LGG subgroups according to the current integrated classification framework (Gue and Lakhani 2024; Louis et al. 2021). Tumors were thus classified into IDH-mutant with 1p/19q codeletion (LGG-IDH-mut-codel, corresponding to oligodendroglioma; TCGA, *n* = 164; CGGA, *n* = 113), IDH-mutant without 1p/19q codeletion (LGG-IDH-mut-non-codel, corresponding to astrocytoma; TCGA, *n* = 235; CGGA, *n* = 142), or IDH-wildtype (LGG-IDH-wt; TCGA, *n* = 91; CGGA, *n* = 88).

**Fig. 1.**
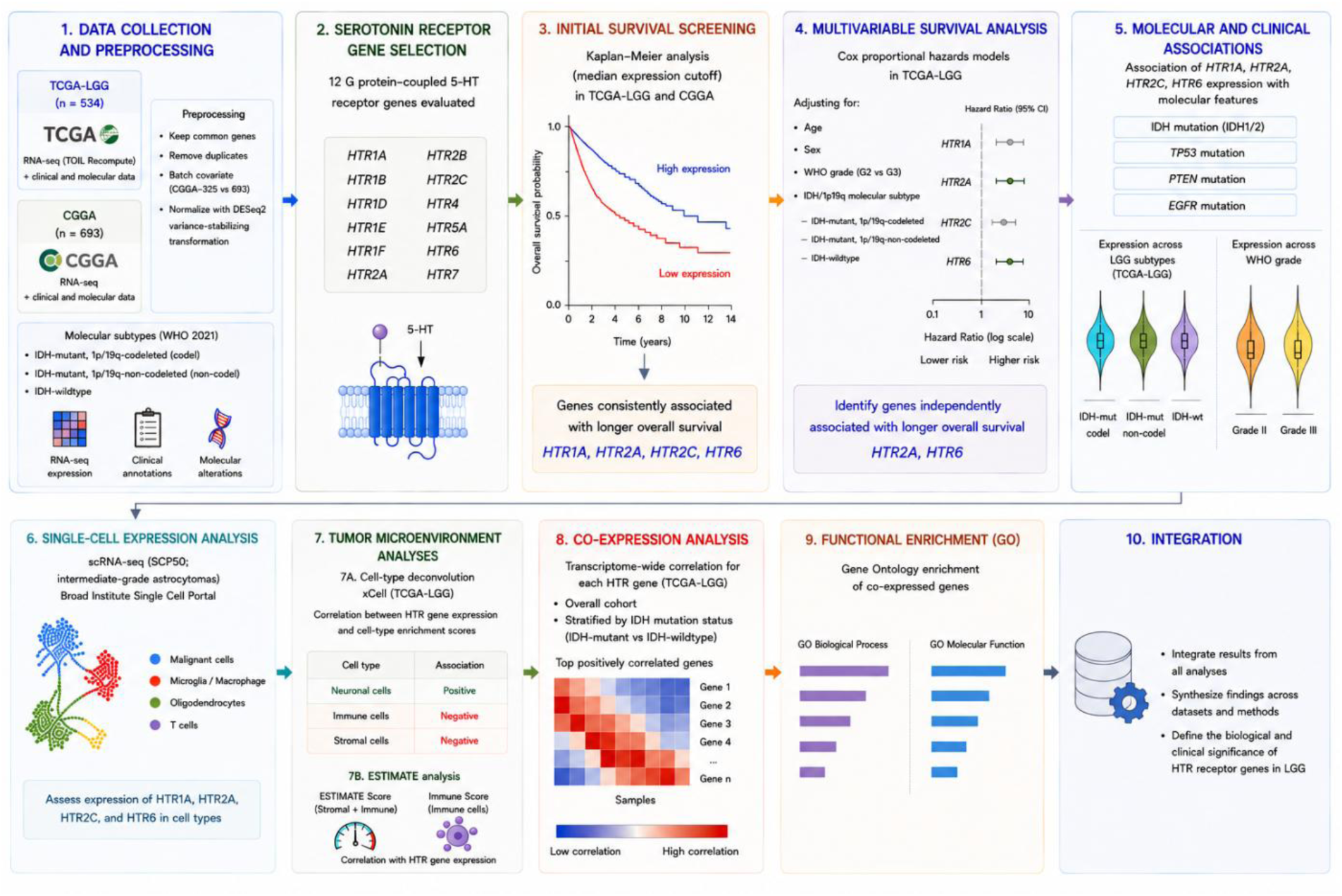
Study workflow and bioinformatic analyses. Schematic overview of the analytical strategy used to investigate G protein-coupled 5-HT receptor genes in LGG. RNA-sequencing, clinical, and molecular data from the TCGA-LGG and CGGA cohorts were used for initial survival screening and subsequent multivariable survival, molecular, and clinical association analyses. Expression of the selected *HTR* genes was further examined at single-cell resolution using the Broad Institute Single Cell Portal SCP50 dataset. Tumor microenvironment associations were assessed using xCell cell-type deconvolution and ESTIMATE scores. Transcriptome-wide co-expression analyses were performed in the overall TCGA-LGG cohort and after stratification by IDH mutation status, followed by Gene Ontology (GO) enrichment analyses for Biological Process and Molecular Function. Results from these complementary analyses were integrated to characterize the biological and clinical significance of 5-HT receptor gene expression in LGG.

For TCGA-LGG, gene-level expression counts generated by the TOIL recompute project were accessed using UCSCXenaTools. Corresponding clinical and molecular information was obtained from the TCGA Pan-Cancer Atlas through cBioPortalData. CGGA data consisted of RNA-sequencing and clinical information from the CGGA-325 and CGGA-693 cohorts, which were downloaded separately and subsequently integrated. Genes not represented in both CGGA datasets were removed before further analysis. To account for technical differences between CGGA-325 and CGGA-693, dataset identity was incorporated as a covariate in analyses performed with DESeq2. Samples for which essential clinical variables or survival information were unavailable were excluded.

Expected counts from TCGA-LGG were converted into raw count estimates before count-based analyses with DESeq2. For CGGA, the count matrices from CGGA-325 and CGGA-693 were merged after gene identifiers had been harmonized. In instances in which more than one transcript mapped to the same gene symbol, the transcript displaying the highest mean expression across samples was selected. Within each cohort, normalized counts were variance-stabilized with DESeq2 and subsequently standardized on a gene-by-gene basis using z-score transformation.

### Survival Analysis

Associations between 5-HT receptor gene expression and overall survival (OS) were initially screened for all 12 active, cloned genes encoding G protein-coupled 5-HT receptors (*HTR1A*, *HTR1B*, *HTR1D*, *HTR1E*, *HTR1F*, *HTR2A*, *HTR2B*, *HTR2C*, *HTR4*, *HTR5A*, *HTR6*, and *HTR7*) in both the TCGA-LGG and CGGA cohorts, using GlioVis data portal for visualization and analysis of brain tumor expression datasets (https://gliovis.bioinfo.cnio.es/; Bowman et al. 2017). This screening identified *HTR1A*, *HTR2A*, *HTR2C*, and *HTR6* as the genes showing the most consistent associations between higher expression and longer OS across both cohorts, and these four genes were selected for subsequent analyses.

The relationships between the expression levels of *HTR1A*, *HTR2A*, *HTR2C*, and *HTR6* and OS were examined using Kaplan–Meier analysis. For each gene, patients were assigned to high- or low-expression groups using the median expression value as the cutoff. Differences in OS between groups were evaluated with the log-rank test. To correct for multiple comparisons, *P*-values obtained across all genes and cohorts were adjusted using the Benjamini–Hochberg (BH) procedure to control the false discovery rate (FDR).

Statistical analyses were conducted in R (version 4.5.1). DESeq2 (version 1.48.1) was used for variance-stabilizing transformation and adjustment for dataset-related effects. Survival analyses were carried out with the survival package (version 3.8.3), using median gene expression to define the patient groups, and Kaplan–Meier plots were generated with survminer. Additional graphical representations were created with ggplot2 (version 4.0.2). UCSCXenaTools (version 1.7.0) and cBioPortalData (version 2.20.0) were used for data retrieval.

### Multivariable Cox Regression

To determine whether 5-HT receptor gene expression was independently associated with OS, separate multivariable Cox proportional hazards regression models were constructed for *HTR1A*, *HTR2A*, *HTR2C*, and *HTR6*. OS (months) and vital status were obtained from the TCGA-LGG clinical annotation. Each model included gene expression together with age at diagnosis, sex, WHO grade, and IDH/1p19q molecular subtype as covariates. Cases with incomplete data for any variable included in the model were excluded from the analysis. Hazard ratios (HRs), 95% confidence intervals (95% CIs), and Wald test *P* values were calculated for each 5-HT receptor. The adjusted hazard ratios of the four receptors were summarized in a forest plot.

### Gene Expression

Gene expression levels in TCGA LGG tumors classified according to subtype, grade, IDH mutations, and oncogene status, were assessed with the GlioVis data portal for visualization and analysis of brain tumor expression datasets, with Tukey’s Honest Significant Difference (HSD) tests used to evaluate differences between groups.

### Single-Cell RNA-Sequencing Data

To determine whether 5-HT receptor genes are expressed by malignant glioma cells, expression of *HTR1A*, *HTR2A*, *HTR2C*, and *HTR6* was examined in a publicly available scRNA-seq dataset of IDH-mutant astrocytoma (SCP50; Broad Institute Single Cell Portal; https://singlecell.broadinstitute.org/single_cell/study/SCP50/single-cell-rna-seq-analysis-of-astrocytoma; Venteicher et al. 2017). Expression patterns were evaluated across the annotated cellular populations provided by the dataset, including malignant cells, oligodendrocytes, microglia/macrophages, and T cells, using the portal’s interactive visualization tools, including t-SNE expression maps and violin plots.

### Deconvolution Analysis

To examine whether 5-HT receptor expression was associated with specific cellular components of the LGG tumor microenvironment (TME), xCell enrichment scores (Aran et al. 2017) were correlated with the expression levels of *HTR1A*, *HTR2A*, *HTR2C*, and *HTR6* in TCGA LGG samples. TCGA expression-matrix sample identifiers were truncated to the first 16 characters to match the sample-level identifiers used in the xCell output. Only samples present in both datasets were retained, and samples were arranged in identical order before analysis.

Pearson correlation coefficients were calculated between the expression of each 5-HT receptor gene and each xCell-derived cell-type or microenvironment enrichment score. Two-sided *P* values were obtained using Pearson correlation tests and corrected across all tested gene–cell-type combinations using the BH FDR method. For visualization, the 15 xCell scores showing the largest absolute Pearson correlation coefficients were selected separately for each receptor. Positive and negative correlations were displayed as horizontal bar plots. Complete correlation coefficients, raw *P* values, and FDR-adjusted *P* values were retained in supplementary tables.

### Analysis of Stromal and Immune Cell Infiltration

Stromal and immune infiltration were estimated using the ESTIMATE algorithm (Yoshihara et al. 2013) implemented in the R package estimate (version 1.0.13). The TCGA-LGG expression matrix was reformatted according to package requirements, and stromal, immune, and ESTIMATE scores were calculated using the estimateScore function. Associations between these scores and 5-HT receptor gene expression were evaluated using the Pearson correlation coefficient (*r*).

### Transcriptome-Wide Co-Expression

Gene-level TPM values (tpm_unstrand assay) were extracted from the SummarizedExperiment object, transformed as log2(TPM + 1), and annotated using HGNC gene symbols. Duplicate gene symbols were removed by retaining the first occurrence. The four 5-HT receptor genes, *HTR1A*, *HTR2A*, *HTR2C*, and *HTR6* were individually evaluated. For each receptor, Pearson correlation coefficients were calculated between its expression levels and those of every other gene across all primary LGG samples. Corresponding *P* values were calculated using cor.test(), and multiple testing correction was performed using the BH FDR procedure. Complete correlation tables were exported for each receptor, and the 30 genes showing the strongest positive correlations were selected for downstream visualization.

Heatmaps were generated using normalized log2(TPM + 1) expression values for each 5-HT receptor together with its 30 strongest positively correlated genes. Samples were hierarchically clustered according to Euclidean distance using complete linkage. Expression values were displayed as row-wise Z scores to facilitate visualization of relative expression patterns across tumors. Clinical annotations, including molecular subtype according to IDH mutation and 1p/19q codeletion status, were incorporated into the heatmaps.

### Gene Ontology

For each 5-HT receptor gene, genes showing a positive Pearson correlation with receptor expression and a BH FDR < 0.05 were ranked according to correlation coefficient. The 200 genes showing the strongest positive correlations with each receptor were selected for Gene Ontology (GO) enrichment analysis. Gene symbols were converted to Entrez Gene identifiers using the bitr() function and the org.Hs.eg.db annotation package. GO enrichment was performed separately for Biological Process (GO:BP) and Molecular Function (GO:MF) categories using the enrichGO() function implemented in the clusterProfiler R package. Statistical significance was assessed using BH correction, with adjusted *P* and *q* value thresholds of 0.05. The 15 highest-ranking enriched terms were displayed for each receptor using dot plots.

### Use of Generative Artificial Intelligence

During preparation of this manuscript, OpenAI’s ChatGPT (GPT-5.5) was used to assist in figure generation and in the identification of grammar and language issues during the final revision of the manuscript. All data analyses, scientific interpretation, table and figure content, and final editing were performed and verified by the authors, who take full responsibility for the published work.

## Results

### Identification of *HTR1A*, *HTR2A*, *HTR2C*, and *HTR6* as 5-HT Receptor Genes Associated with Better Glioma Patient Survival

We initially screened the expression of all 12 cloned genes encoding G protein-coupled 5-HT receptors (*HTR1A*, *HTR1B*, *HTR1D*, *HTR1E*, *HTR1F*, *HTR2A*, *HTR2B*, *HTR2C*, *HTR4*, *HTR5A*, *HTR6*, and *HTR7*), for associations with OS in patients from the TCGA-LGG and CGGA datasets. These early survival analyses were performed using the GlioVis platform (Bowman et al. 2017) with the median expression level as the cutoff. This initial screening identified four genes, *HTR1A*, *HTR2A*, *HTR2C*, and *HTR6,* most consistently associated with longer OS, all four with high expression indicating better survival in both datasets (Fig. 2). These four genes were then selected for subsequent analyses.

**Fig. 2.**
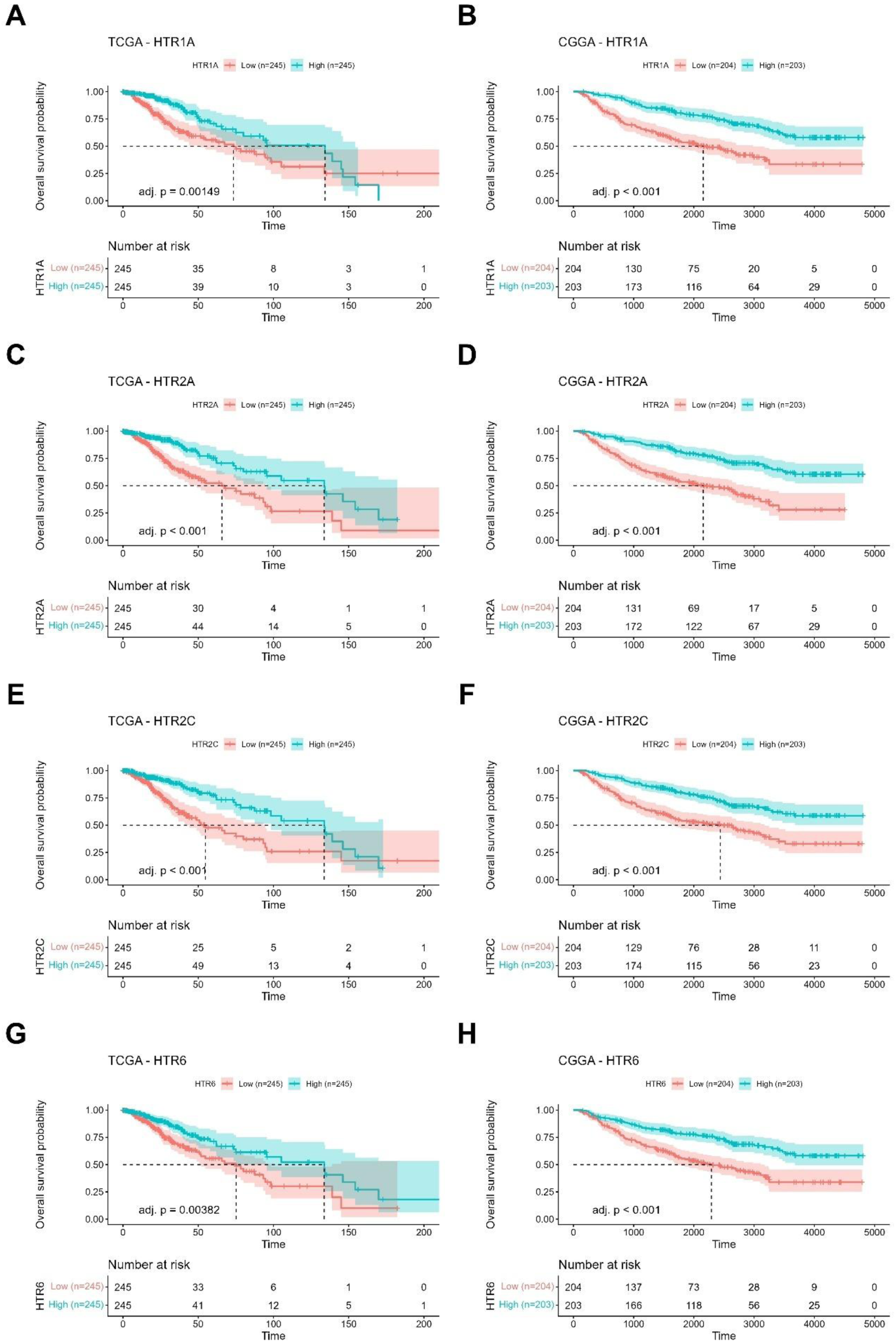
Association between 5-HT receptor gene expression and OS in LGG. Kaplan–Meier survival analyses for *HTR1A* (**A**, **B**), *HTR2A* (**C**, **D**), *HTR2C* (**E**, **F**), and *HTR6* (**G**, **H**) in the TCGA-LGG (**A**, **C**, **E**, **G**) and CGGA (**B**, **D**, **F**, **H**) cohorts. Patients were dichotomized into high- and low-expression groups using median gene expression as the cutoff. Survival distributions were compared using the log-rank test. The number of samples and BH FDR adjusted *P*-values are indicated in the panels.

### *HTR2A* and *HTR6* are Potential Independent Prognostic Factors for Better Survival in Glioma

To determine whether the association between the four 5-HT receptor genes and OS was independently associated with OS, separate multivariable Cox proportional hazards regression models were constructed for *HTR1A*, *HTR2A*, *HTR2C*, and *HTR6*. A total of 450 TCGA patients with complete clinicopathological and transcriptomic data were included in the multivariable Cox regression analyses, of whom 79 experienced the event of interest during follow-up. After adjustment for age, sex, WHO grade, and IDH/1p19q molecular subtype, higher *HTR2A* expression remained independently associated with longer OS (HR = 0.76, 95% CI 0.61–0.94, *P* = 0.011), and an even stronger independent association was observed for *HTR6* (HR = 0.54, 95% CI 0.36–0.81, *P* = 0.003). In contrast, *HTR1A* (HR = 0.84, 95% CI 0.67–1.06, *P* = 0.133) and *HTR2C* (HR = 0.78, 95% CI 0.59–1.03, *P* = 0.084) were not independently associated with survival after adjustment for clinicopathological variables (Fig. 3). These findings indicate that the favorable prognostic value of *HTR2A* and *HTR6* cannot be fully explained by differences in patient age, sex, tumor grade, or IDH/1p19q molecular subtype, supporting the hypothesis that these receptors are linked to biological programs associated with a less aggressive glioma phenotype.

**Fig. 3.**
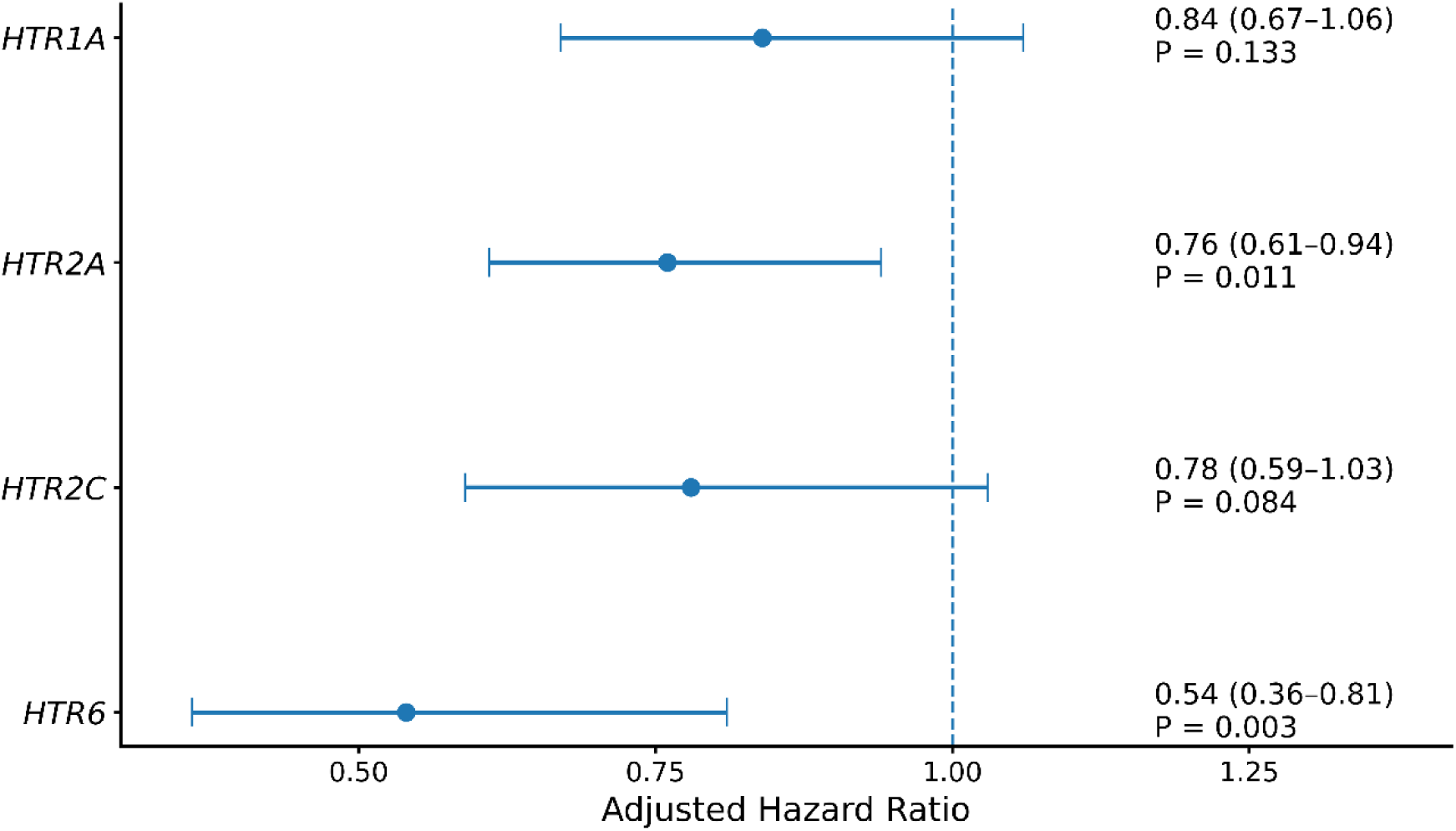
Multivariable associations between 5-HT receptor gene expression and OS in the TCGA-LGG cohort (*n* = 450). Forest plot showing adjusted hazard ratios (HRs) and 95% confidence intervals for *HTR1A*, *HTR2A*, *HTR2C*, and *HTR6* expression. Separate Cox proportional hazards models were adjusted for age, sex, WHO grade, and IDH/1p19q molecular subtype. The dashed vertical line indicates HR = 1.0.

### Expression of *HTR1A*, *HTR2A*, *HTR2C*, and *HTR6* in Glioma According to Subtype and Grade

Comparisons of gene expression across TCGA-LGG molecular subtypes (Louis et al. 2021) showed that *HTR1A* and *HTR2A* were most highly expressed in IDH-mut-codel tumors and least expressed in IDH-wt tumors, with intermediate levels in IDH-mut-non-codel tumors. *HTR2C* expression was also higher in IDH-mut-codel than in IDH-wt tumors, whereas *HTR6* expression was significantly higher in IDH-mut-codel tumors than in both IDH-wt and IDH-mut-non-codel tumors (Fig. 4, Table 1). *HTR1A*, *HTR2A*, and *HTR2C*, but not *HTR6*, were more highly expressed in Grade II than Grade III tumors (Fig. 5, Table 1).

**Fig. 4.**
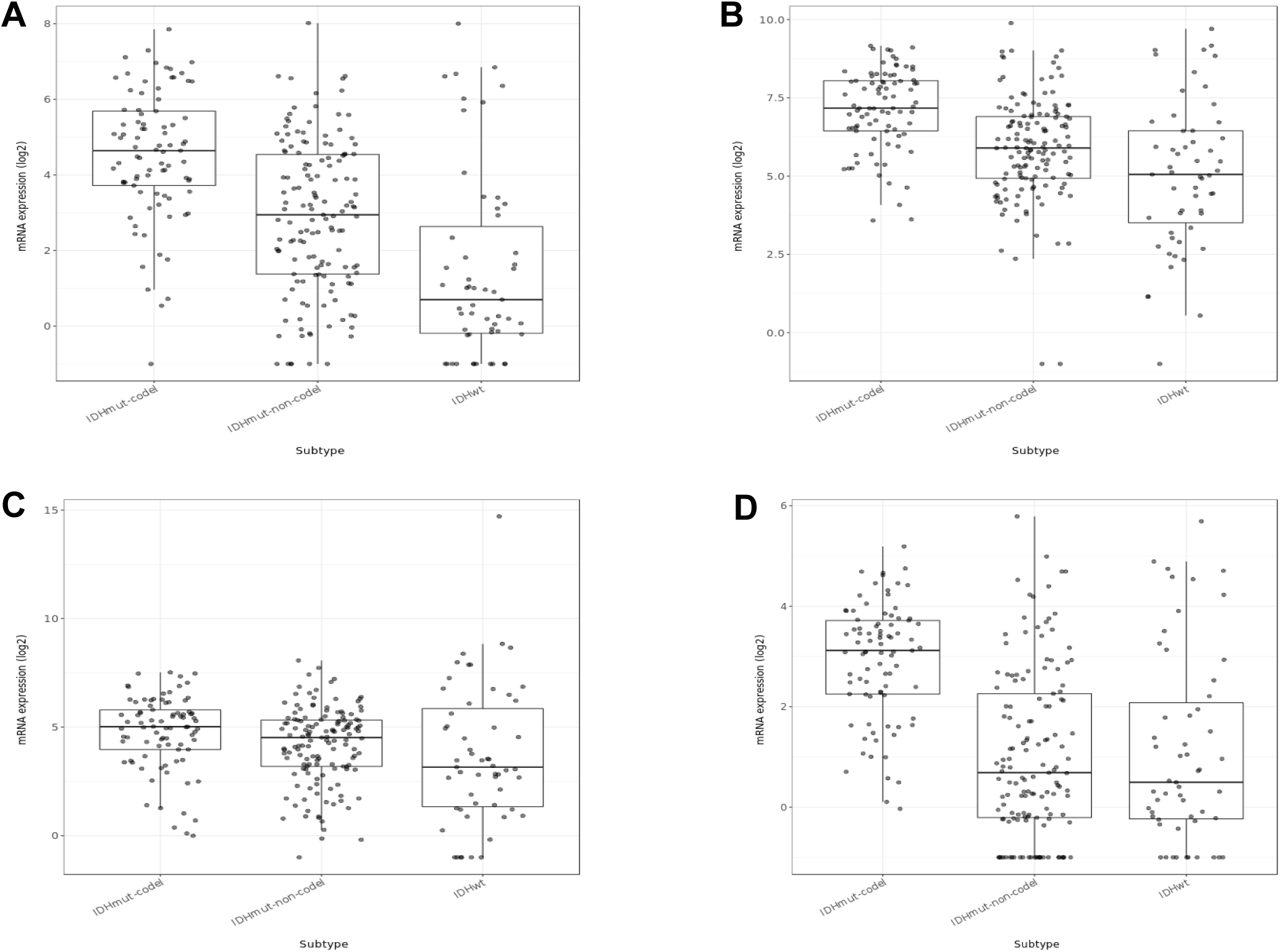
Expression of 5-HT receptor genes according to molecular subtype in TCGA-LGG tumors. Expression levels of *HTR1A* (**A**), *HTR2A* (**B**), *HTR2C* (**C**), and *HTR6* (**D**) in IDH-mut-codel, IDH-mut-non-codel, and IDH-wt tumors; *n* = 281 tumors for which gene expression was matched with subtype annotation in Gliovis (https://gliovis.bioinfo.cnio.es/). Statistical comparison results are summarized in Table 1.

**Fig. 5.**
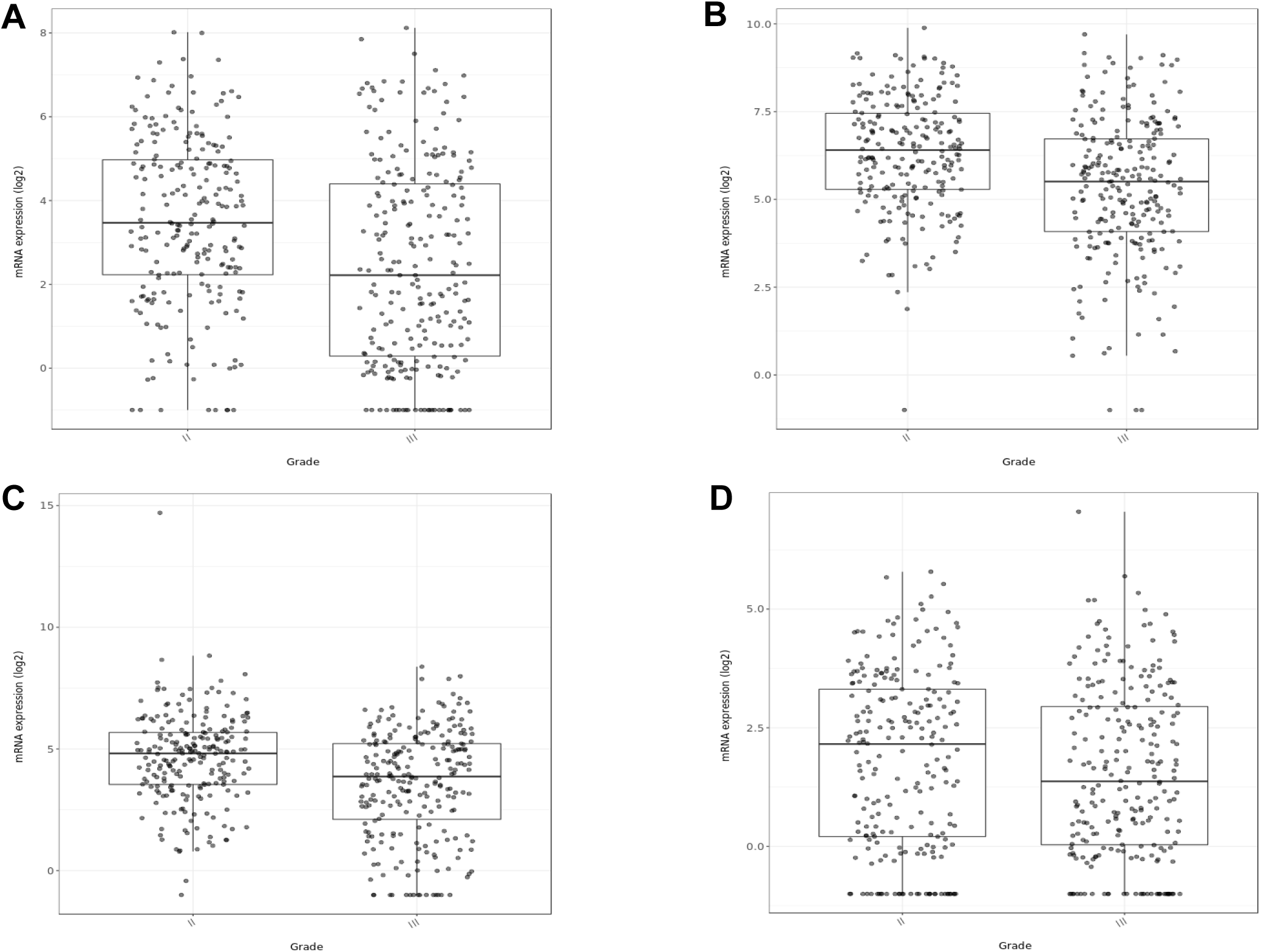
Expression of 5-HT receptor genes according to tumor grade in TCGA-LGG tumors. Levels of *HTR1A* (**A**), *HTR2A* (**B**), *HTR2C* (**C**), and *HTR6* (**D**) in WHO grade II and grade III tumors; *n* = 470 tumors for which gene expression was matched with grade in Gliovis (https://gliovis.bioinfo.cnio.es/). Statistical comparison results are reported in Table 1.

**Table 1.** Statistics for comparisons in expression of *HTR1A*, *HTR2A*, *HTR2C*, and *HTR6* in TCGA LGG tumors across different molecular subtypes and grades.

|  | Diff | Lower | Upper | Adjusted *P*-value | Significance |
| --- | --- | --- | --- | --- | --- |
| ***HTR1A*** |  |  |  |  |  |
| **Tumor subtype** |  |  |  |  |  |
| IDHwt-IDHmut-non-codel | -1.41 | -2.17 | -0.66 | < 0.001 | *** |
| IDHmut-non-codel-IDHmut-codel | -1.65 | -2.30 | -1.00 | < 0.001 | *** |
| IDHwt-IDHmut-codel | -3.06 | -3.89 | -2.24 | < 0.001 | *** |
| **Tumor grade** |  |  |  |  |  |
| II-III | -1.06 | -1.46 | -0.65 | < 0.001 | *** |
| ***HTR2A*** |  |  |  |  |  |
| **Tumor subtype** |  |  |  |  |  |
| IDHwt-IDHmut-non-codel | -0.81 | -1.46 | -0.17 | 0.01 | ** |
| IDHmut-non-codel-IDHmut-codel | -1.26 | -1.82 | -0.70 | < 0.001 | *** |
| IDHwt-IDHmut-codel | -2.07 | -2.78 | -1.37 | < 0.001 | *** |
| **Tumor grade** |  |  |  |  |  |
| II-III | -0.97 | -1.30 | -0.64 | < 0.001 | *** |
| ***HTR2C*** |  |  |  |  |  |
| **Tumor subtype** |  |  |  |  |  |
| IDHwt-IDHmut-non-codel | -0.51 | -1.30 | 0.27 | 0.27 | NS |
| IDHmut-non-codel-IDHmut-codel | -0.59 | -1.26 | 0.09 | 0.11 | NS |
| IDHwt-IDHmut-codel | -1.10 | -1.95 | -0.24 | 0.01 | ** |
| **Tumor grade** |  |  |  |  |  |
| II-III | -1.05 | -1.42 | -0.68 | < 0.001 | *** |
| ***HTR6*** |  |  |  |  |  |
| **Tumor subtype** |  |  |  |  |  |
| IDHwt-IDHmut-non-codel | 0.06 | -0.53 | 0.65 | 0.97 | NS |
| IDHmut-non-codel-IDHmut-codel | -1.82 | -2.46 | -1.17 | < 0.001 | *** |
| IDHwt-IDHmut-codel | -1.88 | -2.38 | -1.37 | < 0.001 | *** |
| **Tumor grade** |  |  |  |  |  |
| II-III | -0.26 | -0.59 | 0.07 | 0.12 | NS |

Diff, estimated difference between the mean expression values of the groups being compared. Lower, lower boundary of the 95% confidence interval for that mean difference. Upper, upper boundary of the 95% confidence interval for that mean difference. NS, non-significant.

### Associations between *HTR1A*, *HTR2A*, *HTR2C*, and *HTR6* Expression and Mutational Status of IDH and Glioma Driver Genes

To further characterize the molecular context associated with 5-HT receptor gene expression, we compared expression levels according to the mutational status of selected key glioma driver genes. *HTR1A* and *HTR2A* expression was significantly higher in tumors harboring *IDH1* mutations, whereas lower expression was observed in tumors with *TP53*, *PTEN*, and *EGFR* mutations. *HTR2C* expression was higher in *IDH2***-**mutant tumors and in tumors with wild-type *PTEN* and *EGFR*, whereas *HTR6* expression was increased in *IDH2*-mutant tumors and in tumors with wild-type *TP53* and *PTEN* (Supplementary Fig. S1-S5, Supplementary Table S1).

Overall, these findings indicate that higher expression of *HTR1A*, *HTR2A*, *HTR2C*, and *HTR6* is preferentially associated with molecular alterations that characterize biologically less aggressive gliomas, particularly *IDH* mutations and absence of alterations in *TP53*, *PTEN*, and *EGFR*, supporting an association between elevated expression of these 5-HT receptor genes and a favorable glioma molecular phenotype.

### Single cell RNA-seq Data Support Malignant Glioma Cells as the Main Source of 5-HT Receptor Gene Expression

An important limitation of using gliomas from datasets such as TCGA and CGGA for transcriptomic studies is that the data is derived from bulk tumor samples, which contain not only tumor cells but also other cell types in the tumor surroundings and TME, such as neurons, microglia, macrophages, oligodendrocytes, and T cell lymphocytes. To address this issue, we analyzed data from the Broad Institute’s Single Cell Portal study SCP50, which contains scRNA seq data for intermediate-grade astrocytomas (Venteicher et al. 2017). The data indicate that expression levels of *HTR1A*, *HTR2A*, *HTR2C*, and *HTR6* are higher in malignant cells compared to other cell types. None of the four genes was detected in oligodendrocytes or T cells, and low levels of only *HTR2A* and *HTR6* are detected in microglia/macrophages (Fig. 6).

**Fig. 6.**
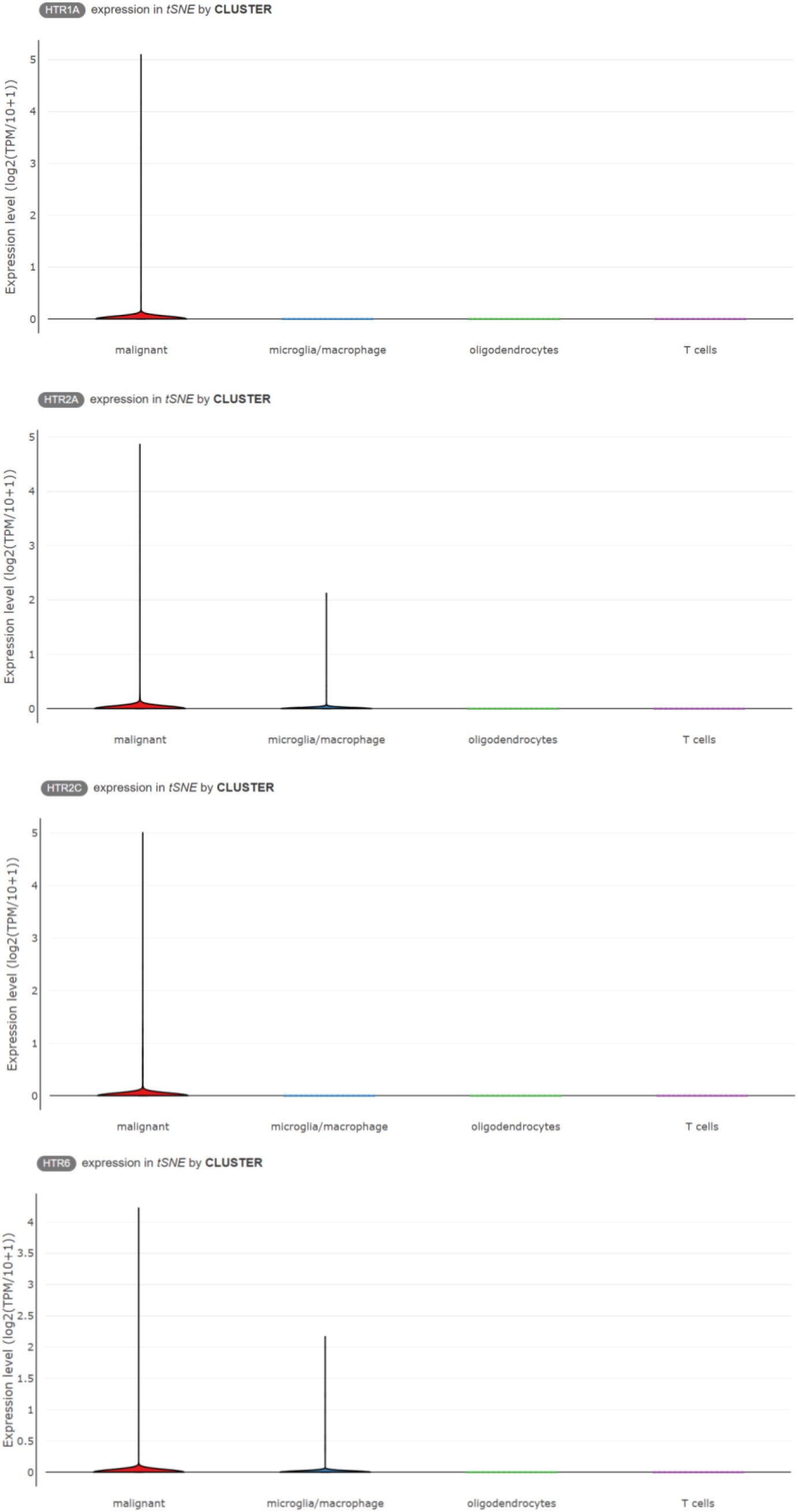
Single-cell expression of 5-HT receptor genes in IDH-mutant astrocytoma. Violin plots showing expression of *HTR1A*, *HTR2A*, *HTR2C*, and *HTR6* across malignant cells, microglia/macrophages, oligodendrocytes, and T cells in the SCP50 single-cell RNA-sequencing dataset. Expression of all four genes was detected predominantly in malignant cells, with lower expression of *HTR2A* and *HTR6* also detected in microglia/macrophages. Malignant cells, *n* = 5,097, microglia/macrophages, *n* = 1,039, oligodendrocytes, *n* = 98, T cells, *n* = 9.

### 5-HT Receptor Gene Expression Is Associated with Neuronal Enrichment and Reduced TME Signatures

Because bulk transcriptomic profiles may also be influenced by differences in cellular composition (Venteicher et al. 2017), we next performed cell-type deconvolution analyses using xCell. Across all four receptor genes, neuronal enrichment represented the strongest positive association among all evaluated signatures (*r* values between 0.55 and 0.80 depending on the receptor). This correlation was particularly pronounced for *HTR1A*, *HTR2A*, and *HTR6*, although *HTR2C* also showed a clear positive association with the neuronal signature. Additional positive correlations varied among individual receptors and included plasmacytoid dendritic cells, regulatory T cells, mesenchymal stem cells, preadipocytes, platelets, and class-switched memory B cells. In contrast, all four receptor genes exhibited negative correlations with multiple indicators of the TME, including the MicroenvironmentScore, StromaScore, ImmuneScore, macrophages, M1 macrophages, mast cells, conventional dendritic cells, endothelial-cell signatures, and Th2 cells (Fig. 7).

**Fig. 7.**
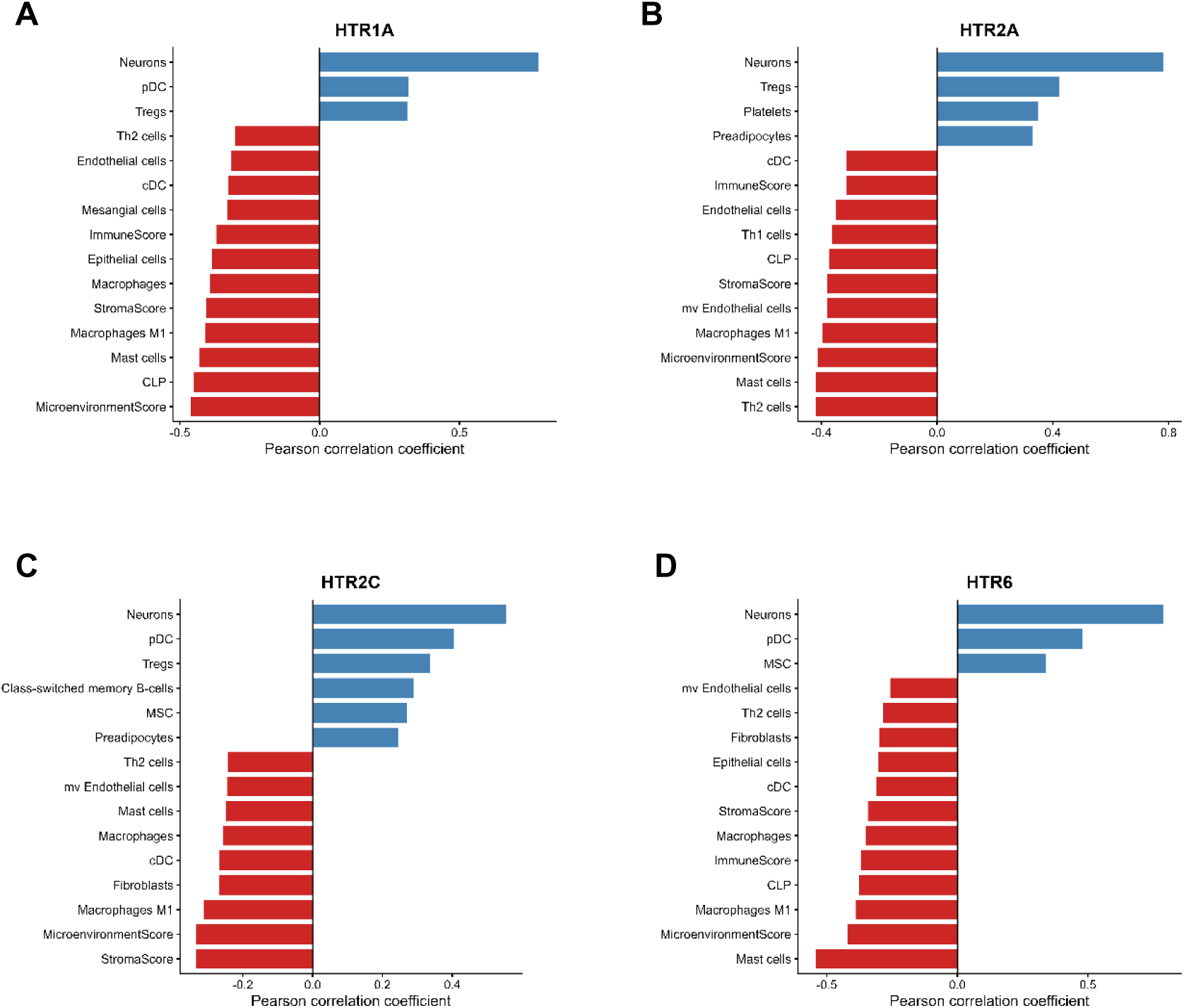
Associations between 5-HT receptor gene expression and cell-type enrichment signatures in TCGA-LGG tumors. Pearson correlations between expression of *HTR1A* (**A**), *HTR2A* (**B**), *HTR2C* (**C**), and *HTR6* (**D**) and xCell-derived enrichment scores (Aran et al. 2017) across 534 TCGA-LGG tumors. The 15 signatures with the largest absolute correlation coefficients for each receptor are shown. Positive and negative correlations are displayed separately.

ESTIMATE analysis demonstrated significant negative correlations between expression of *HTR1A*, *HTR2A*, *HTR2C*, and *HTR6* and all three derived scores (stromal, immune, and ESTIMATE), indicating that tumors with high expression of these genes were associated with lower predicted stromal and immune infiltration (Fig. 8). *HTR1A* and *HTR6* showed the strongest negative correlations, particularly with ImmuneScore (*HTR1A*, *r* = −0.565, FDR = 2.73 × 10⁻⁴⁵; *HTR6*, *r* = −0.558, FDR = 3.06 × 10⁻⁴⁴). *HTR2A* demonstrated intermediate inverse correlations (ImmuneScore, *r* = −0.475, FDR = 4.98 × 10⁻³¹), whereas *HTR2C* displayed weaker but still highly significant associations (ImmuneScore, *r* = −0.355, FDR = 3.21 × 10⁻¹⁷). Similar patterns were observed for StromalScore and ESTIMATEScore. All twelve receptor–score associations remained statistically significant following FDR correction.

**Fig. 8.**
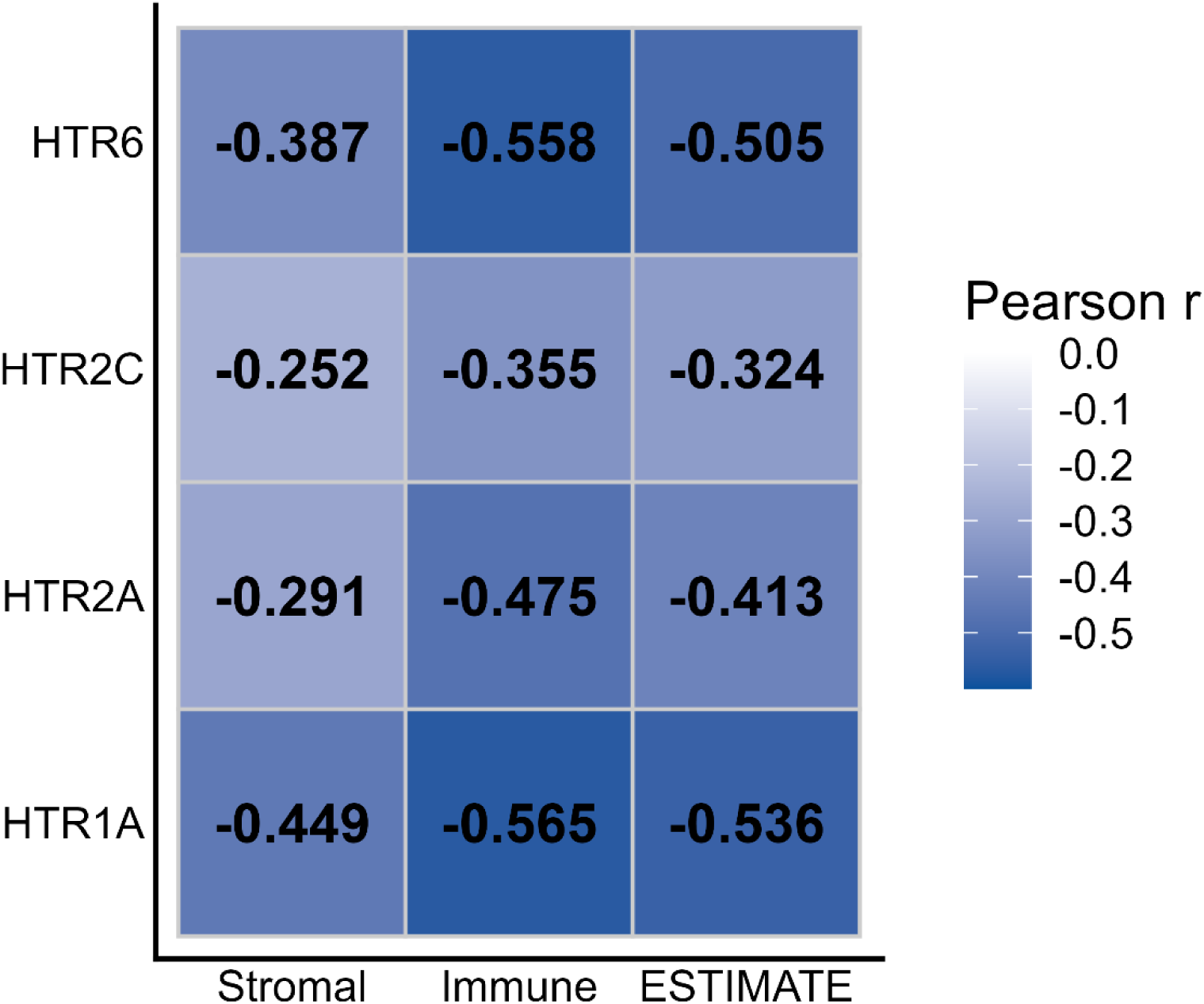
5-HT receptor gene expression is inversely associated with stromal and immune infiltration in TCGA-LGG tumors. Heatmap showing correlations between *HTR1A*, *HTR2A*, *HTR2C*, and *HTR6* expression and StromalScore, ImmuneScore, and ESTIMATEScore derived using ESTIMATE across 534 tumors. Values within cells indicate Pearson correlation coefficients (*r*). All receptor–score associations remained significant after FDR correction.

Together, these results support the view that elevated expression of *HTR1A*, *HTR2A*, *HTR2C*, and *HTR6* characterizes gliomas with a neuronal-like transcriptional program, reduced stromal and immune cell infiltration, and expression occurring predominantly in malignant glioma cells in the single-cell dataset.

### Genome-Wide Transcriptomic Analysis Reveals Positive Correlations between 5-HT Receptor Genes and Genes Involved in Synaptic Function

Transcriptome-wide co-expression analysis demonstrated that each of the four 5-HT receptor genes was associated with a distinct yet partially overlapping neuronal transcriptional program (Fig. 9). *HTR1A* showed strong positive correlations with multiple genes involved in synaptic vesicle function, neurotransmitter release, neuronal differentiation, and excitatory neurotransmission, including *SRRM4*, *STXBP1*, *TAFA2*, *MYT1L*, *RIMS1*, *RIMS2*, *BSN*, *SCN8A*, *KCNT1*, and *GRIN1*. *HTR2A* exhibited a pronounced association with genes related to inhibitory synaptic signaling, including *GABRA1*, *GABRA4*, *GABRB2*, *GABRG2*, *ATP8A2*, *DLGAP2*, *SLC12A5*, *HCN1*, and *ARHGAP44*. *HTR2C*, in turn, also demonstrated a neuronal co-expression profile but with a partially distinct repertoire of correlated genes, including *ATP8A2*, *ARHGAP44*, *DRD5*, *CHRM2*, *EPHA8*, *KCNH5*, and *OCA2*. Finally, *HTR6* displayed one of the strongest neuronal signatures observed, showing robust correlations with genes involved in synaptic transmission and neuronal maturation, including *MYT1L*, *STXBP1*, *DLG4*, *CHRM1*, *KCNC1*, *SCN2A*, *GABRG2*, *SPRN*, *GRM2*, and *RIMS1*. Heatmap analysis indicated that the expression patterns of the 5-HT receptor genes and their top correlated genes were coordinated across glioma samples, consistent with tightly regulated transcriptional programs.

**Fig. 9.**
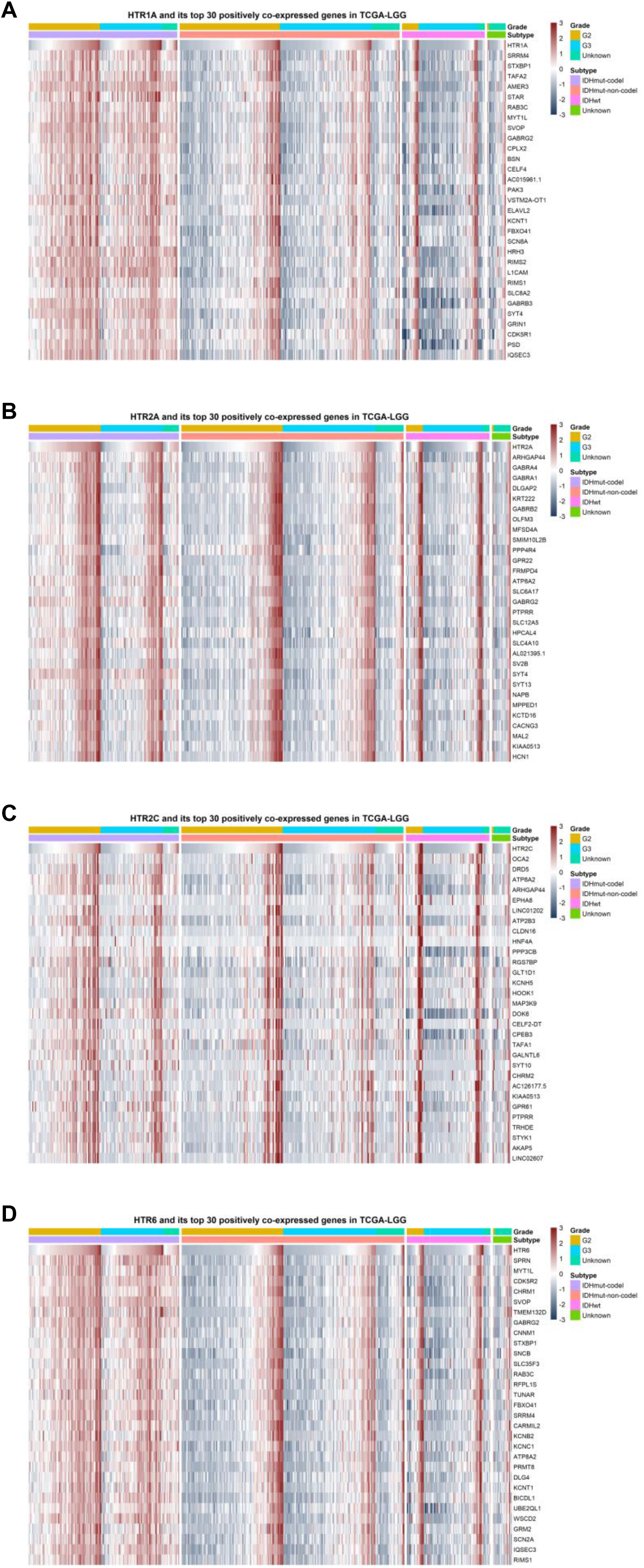
Transcriptome-wide co-expression patterns associated with 5-HT receptor genes in TCGA-LGG tumors. Heatmaps showing *HTR1A* (**A**), *HTR2A* (**B**), *HTR2C* (**C**), and *HTR6* (**D**) together with their 30 strongest positively correlated genes across 534 tumors. Expression values are shown as row-wise Z scores. Samples were hierarchically clustered, with tumor grade and IDH/1p19q molecular subtype indicated by annotation bars.

### Gene Ontology Enrichment Reveals Convergent Synaptic Transcriptional Programs

GO Biological Process enrichment analysis (Fig. 10) showed that genes positively correlated with *HTR1A* expression were strongly enriched for pathways related to neuronal communication and synaptic function. Among the highest-ranking biological processes were regulation of membrane potential, exocytosis, regulation of synaptic plasticity, neurotransmitter transport, synaptic vesicle cycle, neurotransmitter secretion, signal release from synapse, vesicle-mediated transport in synapse, and calcium ion-regulated exocytosis. GO Molecular Function analysis (Fig. 11) similarly demonstrated enrichment of functions associated with neurotransmission, ion channel regulation, receptor activity, and synaptic signaling. Comparable enrichment profiles were observed for *HTR2A*, *HTR2C*, and *HTR6*, indicating that despite belonging to different 5-HT receptor families, all four receptors are involved in similar neuronal and synaptic transcriptional programs in LGG.

**Fig. 10.**
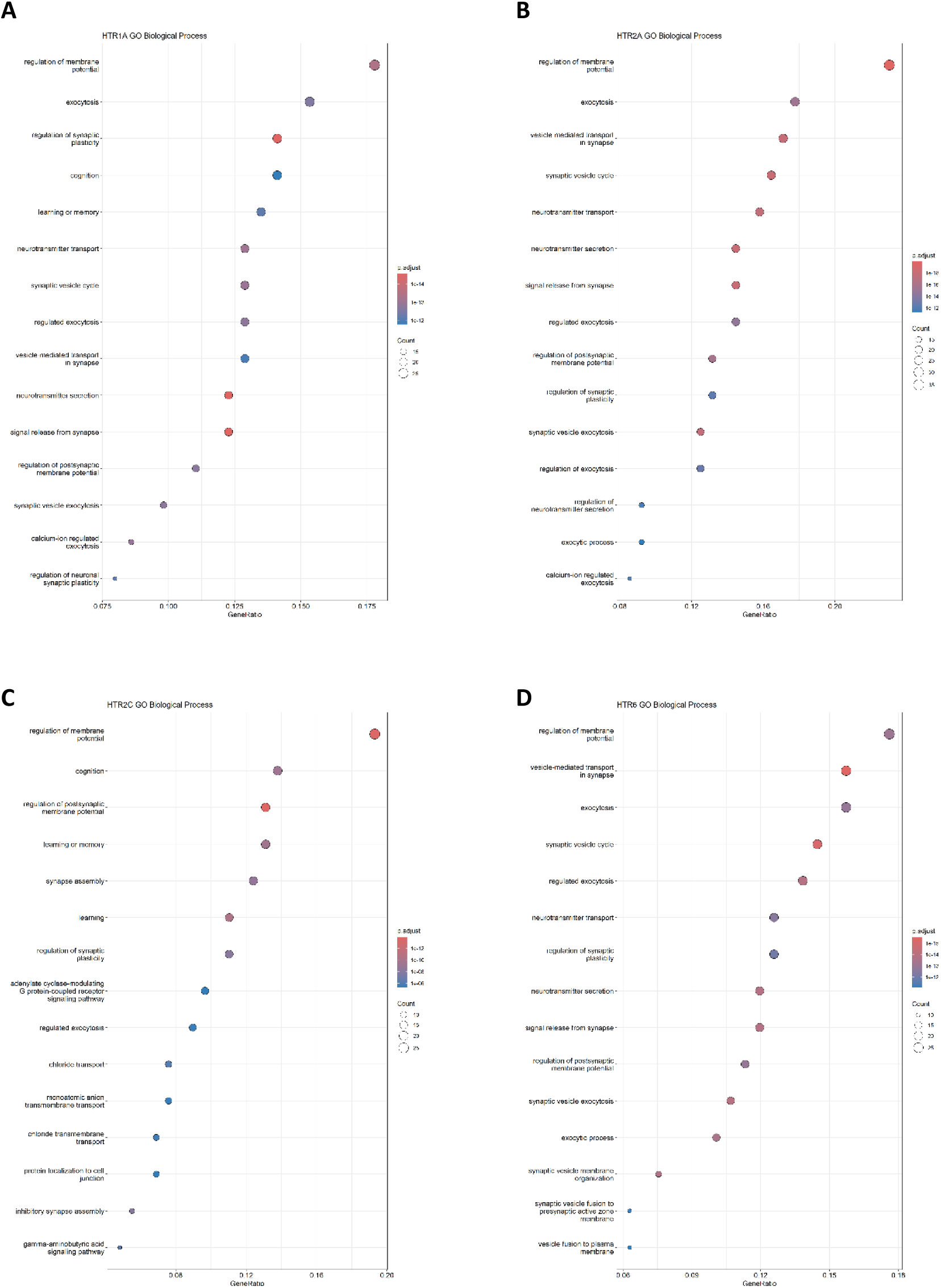
Gene Ontology Biological Process enrichment of genes positively associated with 5-HT receptor expression in LGG. Analysis of the 200 genes showing the strongest positive correlations with genes positively correlated with *HTR1A* (**A**), *HTR2A* (**B**), *HTR2C* (**C**), and *HTR6* (**D**) in 534 TCGA-LGG tumors. Dot size represents the number of genes associated with each term, and color indicates statistical significance. Enriched processes converge predominantly on neuronal communication, membrane excitability, synaptic vesicle trafficking, neurotransmitter release, and synaptic plasticity.

**Fig. 11.**
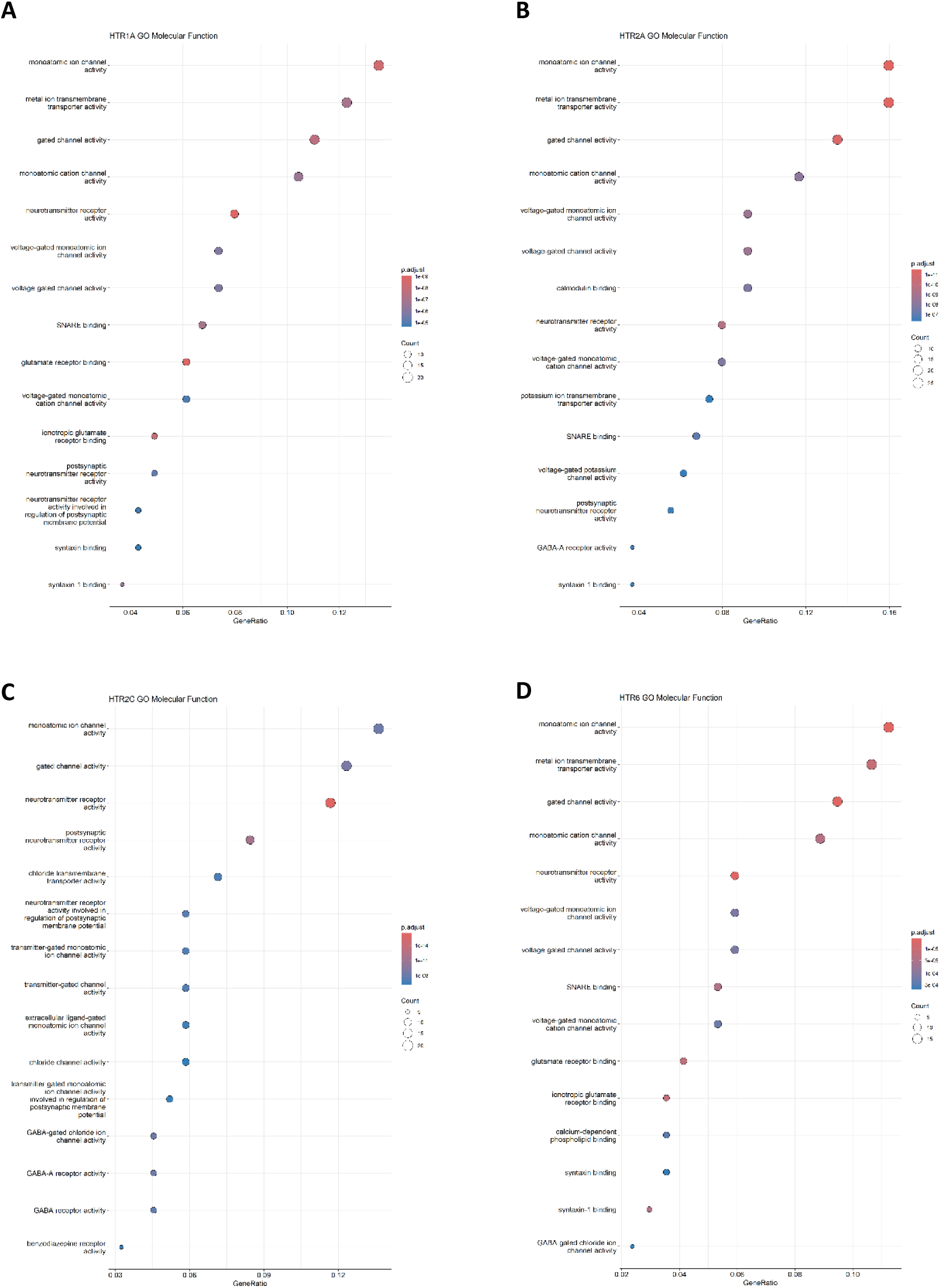
Gene Ontology Molecular Function enrichment of genes positively associated with 5-HT receptor expression in LGG. Analysis of the 200 genes showing the strongest positive correlations with *HTR1A* (**A**), *HTR2A* (**B**), *HTR2C* (**C**), and *HTR6* (**D**) in 534 TCGA-LGG tumors. Dot size represents the number of genes associated with each term, and color indicates statistical significance. Prominent enriched functions include ion channel activity, neurotransmitter receptor activity, transporter activity, and other functions related to neuronal and synaptic signaling.

### Transcriptome-Wide Co-Expression Patterns Differ between IDH-Mutant and IDH-Wild Type Gliomas

Given that IDH mutation status is a major molecular determinant of glioma biology and patient prognosis, transcriptome-wide co-expression analyses were repeated separately in IDH-mutant and IDH-wildtype tumors to determine whether the 5-HT receptor-associated transcriptional programs were preserved within each molecular subgroup. Heatmaps generated from the top positively correlated genes revealed marked differences between IDH-mutant and IDH-wildtype tumors while indicating that the association of 5-HT receptor genes with neuronal transcriptional programs was preserved within both molecular subgroups (Fig. 12). Among IDH-mutant gliomas, all four 5-HT receptor genes showed remarkably consistent co-expression patterns. Although the identity of individual highly correlated genes varied among receptors, the overall biological signatures were conserved and were enriched for genes involved in neuronal differentiation, synaptic organization, synaptic vesicle trafficking, neurotransmitter release, ion channel function, and neuronal signaling. Recurrently identified genes included components of the presynaptic machinery (*RIMS1*, *STXBP1*, *SNAP91*, *SVOP*, and *RAB3C*), neuronal ion channels (*KCNC1* and *SCN8A*), γ-aminobutyric acid (GABA) receptor subunits, and neuronal differentiation markers such as *MYT1L*. These coherent transcriptional modules indicate that high expression of *HTR1A*, *HTR2A*, *HTR2C*, and *HTR6* may identify a shared neuronal/synaptic differentiation program within IDH-mutant gliomas. Among the four receptor genes, *HTR6* exhibited the most pronounced neuronal signature. Its co-expression network showed strong enrichment for canonical neuronal and synaptic genes, including *DLG4*, *SYN1*, *SYP*, *STX1B*, *KCNC1*, *RIMS1*, *GABRG2*, and *MYT1L*, forming one of the most coherent neuronal transcriptional modules observed in the study.

**Fig. 12.**
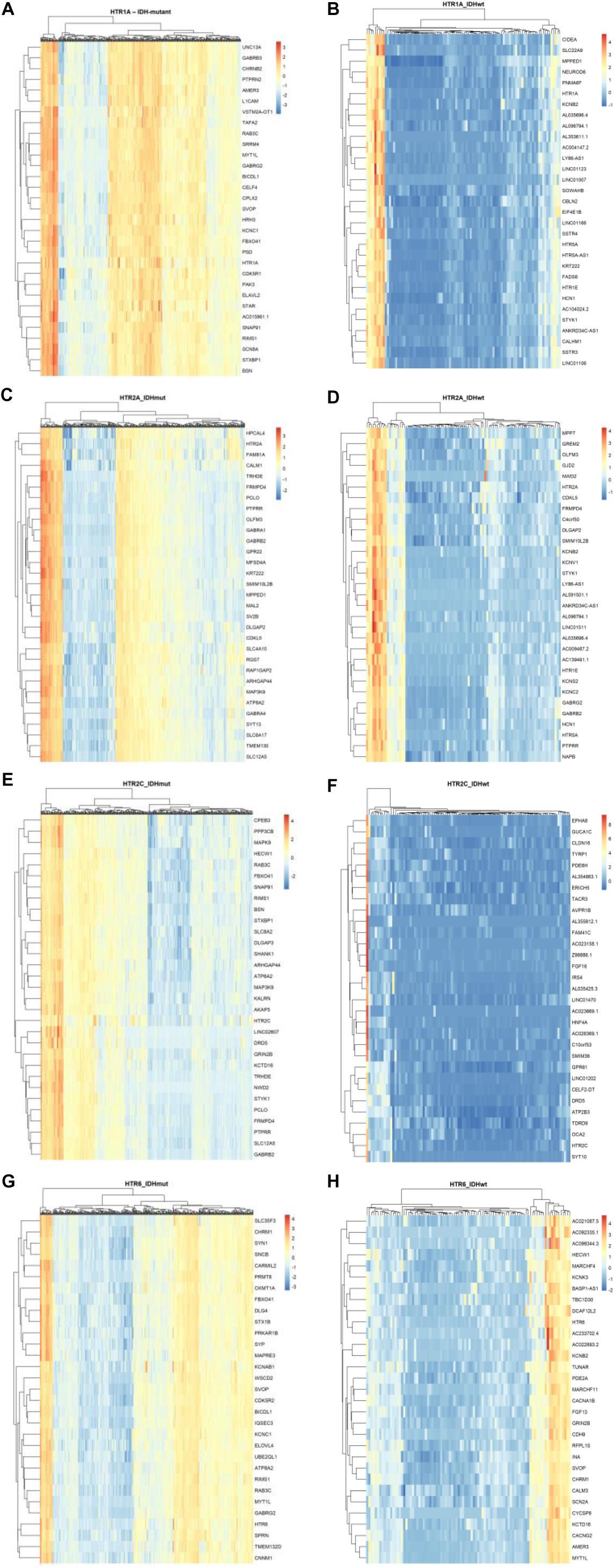
5-HT receptor-associated co-expression programs in IDH-mutant and IDH-wildtype gliomas. Heatmaps showing the top positively correlated genes for *HTR1A* (**A**, **B**), *HTR2A* (**C**, **D**), *HTR2C* (**E**, **F**), and *HTR6* (**G**, **H**), analyzed separately in IDH-mutant (**A**, **C**, **E**, **G**) and IDH-wildtype (**B**, **D**, **F**, **H**) TCGA-LGG tumors. IDH-mutant tumors (*n* = 419) displayed highly coherent neuronal/synaptic co-expression programs across the four receptor genes, whereas IDH-wildtype tumors (*n* = 94) exhibited more heterogeneous transcriptional patterns.

In contrast, IDH-wildtype tumors displayed substantially different co-expression landscapes. Although neuronal genes remained among the positively correlated transcripts for all four receptors, the associated networks were more heterogeneous, less uniformly organized, and contained distinct sets of correlated genes compared with the IDH-mutant subgroup. Receptor genes remained associated with neuronal signaling molecules, ion channels, and neurotransmitter receptor genes, but the characteristic synaptic module observed in IDH-mutant tumors was less prominent. Consistent with this observation, the clustering patterns of IDH-wildtype tumors were more fragmented and showed greater transcriptional heterogeneity than those observed in IDH-mutant gliomas.

Despite these subtype-specific differences, several neuronal genes repeatedly appeared among the top correlated transcripts across receptors and molecular subgroups, including *RIMS1*, *STXBP1*, *KCNC1*, *RAB3C*, *FBXO41*, *MYT1L*, *SNAP91*, *DLG4*, and *ATP8A2*, supporting the existence of a conserved neuronal transcriptional core associated with 5-HT receptor expression in LGG.

## Discussion

*HTR1A* was among the first 5-HT receptor genes to be molecularly identified and was cloned in 1987 as a single intronless gene and identified as a GPCR-coding gene based on sequence homology to the β2-adrenergic receptor gene (Kobilka et al. 1987). *HTR2C* was subsequently cloned and its corresponding protein initially called the 5-HT1C receptor based on its pharmacology (Julius et al. 1988). *HTR2A* soon followed, cloned from a rat brain cDNA library (Pritchett et al. 1988). *HTR6* represents a second generation and was one of the last 5-HT GPCR genes to be discovered, cloned from rat striatal mRNA as a cDNA sequence exhibiting high homology to previously known 5-HT receptor genes (Monsma et al. 1993). These four 5-HT receptor types regulate neuronal excitability, synaptic transmission, and synaptic plasticity through distinct intracellular signaling pathways. 5-HT1A receptors couple predominantly to Gi/o proteins, whereas 5-HT2A and 5-HT2C receptors couple to Gq/11 proteins and 5-HT6 couples to Gs proteins, thereby differentially modulating cAMP- and phospholipase C-dependent signaling in neurons (Barnes et al. 2021). Variants in each of these four 5-HT receptor genes have been implicated in psychiatric disorders and Alzheimer’s disease (Grubor et al. 2020; Holmes et al. 1998; 2003; Illi et al. 2009; Pritchard et al. 2008; Saeb et al. 2026; Thome et al. 2001), and at least *HTR6* has been also implicated in Parkinson’s disease (Messina et al. 2002).

In our study, *HTR2A* and *HTR6* stood out as candidate genes that may be independent prognostic factors for better survival in patients with LGG. In a large breast cancer study, expression of *HTR1A*, *HTR2C*, and *HTR6*, but not *HTR2A*, was positively associated with better relapse-free survival (RFS) rates (Zhang et al. 2022). Other findings indicated that *HTR1A*, *HTR2A*, and *HTR2C*, among other 5-HT receptor genes, were downregulated in breast tumors and associated with RFS in breast cancer patients, whereas *HTR6* was among the genes associated with shorter OS (Zhan et al. 2023). Shen et al. (2023) examined a three-gene set of genes linked to schizophrenia, including *HTR2A*, in a pan-cancer analysis that included Cox regression and patient survival analyses and identified GBM and LGG as tumor types particularly worth of further investigation. *HTR2A* was associated with immune infiltration, tumor mutational burden, and mutant-allele tumor heterogeneity in these glioma types. However, a negative correlation was found between the *HTR2A* upregulation and survival.

Our results also showed that the 5-HT receptor genes are more highly expressed in IDH-mutant than IDH wild-type and in grade II than grade III tumors, and, importantly, *HTR2C* and *HTR6* were more expressed in LGG tumors with no mutations in key glioma driver genes. Although we cannot rule out the possibility that non-tumor cells within the tumor samples, including neurons, contribute to the observed 5-HT receptor gene expression levels, our results from available single-cell data analysis, negative correlations found with TME cell types, and cell type deconvolution findings do not support non-tumor cells as the main source of gene expression. Moreover, 5-HT receptor gene expression was associated with a broader set of genes known to crucially regulate synaptic development and transmission, a pattern that was markedly more pronounced in IDH-mutant than IDH-wild type LGG.

A recent compelling mechanistic study by Michelle Monje and colleagues focused specifically on serotonergic influences on neuronal interactions with high-grade gliomas. Consistent with the view that neuronal activity promotes GBM progression, serotonergic neuron activity stimulated circuit-specific increases in tumor proliferation and was associated with reduced survival. Knockout or pharmacological inhibition of 5HT2A receptors in glioma cells suppressed the growth-promoting effects of 5-HT. The results indicated that GBMs influence serotonergic neuronal activity patterns, leading to 5-HT release into the TME and 5HT2A-mediated GBM stimulation. Moreover, the study reveals a novel feed-forward functional interaction mechanism between serotonergic neurons and GBM (Drexler et al. 2025b).

In LGG, in contrast, our previous transcriptomic analyses of tumors in the TCGA LGG and CGGA cohorts suggested a pattern where higher expression of genes associated with excitatory synaptic activity, including those encoding AMPAR subunits, AMPAR auxiliary proteins, GABA_A_ receptor subunits, postsynaptic scaffolding proteins, and mitochondrial protein-coding genes, was associated with a more favorable tumor biology and a broader neuronal/synaptic gene expression program. These findings raised the hypothesis that LGG tumors showing high expression of synaptic and neuronal genes might display features of a more differentiated and less aggressive cellular phenotype (Badalotti et al. 2024; Dal-Pizzol et al. 2026; Gaia et al. 2026; Rodrigues et al. 2026).

Other evidence supports the view that increased differentiation is associated with less aggressive biology in grade 2 and grade 3 gliomas. High gene expression of the neuronal marker *MAP2* is associated with longer survival in TCGA LGG, and reprogramming glioma cells towards a neuronal-like state with transcription factors SOX11 or ZIC1 suppresses cell proliferation (Fu et al. 2019). Consistent with these observations, experimental reprogramming of grade 3 astrocytoma and oligodendroglioma cells toward neuronal differentiation using small molecules or neural transcription factors reduced proliferation and tumor-associated gene expression while activating tumor-suppressive programs (Yi et al. 2024). Single-cell RNA-seq profiling of *IDH1*- or *IDH2*-mutant oligodendrogliomas identified an undifferentiated, subpopulation of cells marked by a neural stem cell expression program which may drive tumor growth (Tirosh et al. 2016). A recent study integrating single-nucleus transcriptomic profiling with bulk DNA and RNA sequencing of IDH-mutant oligodendrogliomas and astrocytomas from 35 patients showed that malignant cell states transcriptionally resemble stages of normal glial and neuronal lineage development or adopt a mesenchymal-like state similar to those observed in IDH-wild-type GBM. Importantly, less differentiated malignant glioma cell states became more prevalent with increasing tumor grade and the acquisition of oncogenic alterations, while reduced lineage differentiation was associated with genetic features linked to recurrence (Johnson et al. 2026).

The present results are consistent with this emerging model, in which the retention or acquisition of a more differentiated neuronal-like transcriptional state may be associated with less aggressive glioma biology. Increased levels of specific 5-HT receptor types in LGG tumors may be linked to a neural development and synaptic function signature associated with biological features of less malignant tumors and better patient survival. However, this working model is still mostly based solely on correlational transcriptomic analysis of bulk tumor samples, and further functional studies exploring cellular mechanisms and identifying intra-tumoral cell types are required.

As our previous reports cited above, the present study has a number of important limitations. First, the biological roles of 5-HT receptor genes in LGG were inferred from transcriptomic associations and therefore do not establish causal relationships. Future studies integrating functional experiments, single-cell transcriptomics, spatial transcriptomics, and multi-omic strategies will be required to determine whether the 5-HT receptor-associated transcriptional programs identified here actively contribute to tumor biology or reflect differences in cellular differentiation states. Second, although we have included limited single-cell data and cell-type deconvolution analysis, given that the study was based primarily on bulk tumor transcriptomes, the precise cellular origin of the observed gene expression patterns still cannot be established with certainty.

## Conclusion

Our findings identify *HTR1A*, *HTR2A*, *HTR2C*, and *HTR6* as a subset of 5-HT GPCR genes associated with favorable clinical and molecular features in LGG, with *HTR2A* and *HTR6* showing independent associations with longer survival. Across complementary bulk transcriptomic, single-cell, deconvolution, and functional enrichment analyses, expression of these receptors consistently marked a neuronal and synaptic transcriptional state characterized by genes involved in synaptic organization, neurotransmitter release, membrane excitability, and neuronal differentiation. This program was particularly coherent in IDH-mutant gliomas, suggesting that 5-HT receptor expression may constitute part of a broader neuronal differentiation state associated with less aggressive tumor biology. Functional studies will be required to determine whether these receptors actively contribute to this phenotype or primarily serve as markers of the underlying transcriptional state.

## Author Contributions

Conceptualization, F.L.C.F., G.R.I., R.R.; methodology, F.L.C.F., H.R.D.P., G.R.I., R.R.; formal analysis, F.L.C.F., H.R.D.P., R.R.; investigation, F.L.C.F., H.R.D.P., R.R; data curation, F.L.C.F., H.R.D.P., R.R.; writing – original draft, R.R.; Writing – review & editing, F.L.C.F., H.R.D.P., G.R.I., R.R.; Supervision, G.R.I., R.R.; Project administration, G.R.I., R.R.; Funding acquisition, G.R.I., R.R.

## Data Availability

The original datasets used in the study are openly available in The Cancer Genome Atlas (TCGA) Research Network (https://www.cancer.gov/tcga), the Chinese Glioma Genome Atlas (CGGA, http://www.cgga.org.cn), and the Broad Institute Single Cell Portal (https://singlecell.broadinstitute.org/single_cell/study/SCP50/single-cell-rna-seq-analysis-of-astrocytoma). The R scripts used for data processing, survival analyses, transcriptome-wide co-expression analysis, Gene Ontology enrichment, tumor microenvironment analyses, and figure generation in this study are publicly available at GitHub: https://github.com/rafaelroesler/HTR-TCGA-LGG.

## Declarations

### Consent to Participate

Not applicable.

### Human Ethics and Consent to Participate

This study was based exclusively on analyses of publicly available, de-identified transcriptomic datasets and publicly accessible online bioinformatic resources. No human participants were recruited, no human tissue or biological specimens were collected or analyzed, no identifiable patient information was accessed, and no animal experiments were performed. Therefore, institutional ethics committee approval and informed consent were not required.

## References

1. Aran D, Hu Z, Butte AJ (2017) xCell: digitally portraying the tissue cellular heterogeneity landscape. Genome Biol 18(1): 220. doi: 10.1186/s13059-017-1349-1

2. Badalotti R, Dalmolin M, Malafaia O, Ribas Filho JM, Roesler R, Fernandes MAC, Isolan GR (2024) Gene expression of GABA_A_ receptor subunits and association with patient survival in glioma. Brain Sci 14(3): 275. doi: 10.3390/brainsci14030275

3. Barnes NM, Ahern GP, Becamel C, Bockaert J, Camilleri M, Chaumont-Dubel S, Claeysen S, Cunningham KA, Fone KC, Gershon M, Di Giovanni G, Goodfellow NM, Halberstadt AL, Hartley RM, Hassaine G, Herrick-Davis K, Hovius R, Lacivita E, Lambe EK, Leopoldo M, Levy FO, Lummis SCR, Marin P, Maroteaux L, McCreary AC, Nelson DL, Neumaier JF, Newman-Tancredi A, Nury H, Roberts A, Roth BL, Roumier A, Sanger GJ, Teitler M, Sharp T, Villalón CM, Vogel H, Watts SW, Hoyer D (2021) International Union of Basic and Clinical Pharmacology. CX. Classification of Receptors for 5-hydroxytryptamine; Pharmacology and Function. Pharmacol Rev 73(1): 310–520. doi: 10.1124/pr.118.015552

4. Barron T, Yalçın B, Su M, Byun YG, Gavish A, Shamardani K, Xu H, Ni L, Soni N, Mehta V, Maleki Jahan S, Kim YS, Taylor KR, Keough MB, Quezada MA, Geraghty AC, Mancusi R, Vo LT, Castañeda EH, Woo PJ, Petritsch CK, Vogel H, Kaila K, Monje M (2025) GABAergic neuron-to-glioma synapses in diffuse midline gliomas. Nature 639(8056): 1060–1068. doi: 10.1038/s41586-024-08579-3

5. Bockaert J, Claeysen S, Bécamel C, Dumuis A, Marin P (2006) Neuronal 5-HT metabotropic receptors: fine-tuning of their structure, signaling, and roles in synaptic modulation. Cell Tissue Res 326(2): 553–572. doi: 10.1007/s00441-006-0286-1

6. Bowman RL, Wang Q, Carro A, Verhaak RG, Squatrito M (2017) GlioVis data portal for visualization and analysis of brain tumor expression datasets. Neuro Oncol 19(1): 139–141. doi: 10.1093/neuonc/now247

7. Bready D, Placantonakis DG (2019) Molecular pathogenesis of low-grade glioma. Neurosurg Clin N Am 30(1): 17–25. doi: 10.1016/j.nec.2018.08.011

8. Cancer Genome Atlas Research Network; Brat DJ, Verhaak RG, Aldape KD, Yung WK, Salama SR, Cooper LA, Rheinbay E, Miller CR, Vitucci M, Morozova O, Robertson AG, Noushmehr H, Laird PW, Cherniack AD, Akbani R, Huse JT, Ciriello G, Poisson LM, Barnholtz-Sloan JS, Berger MS, Brennan C, Colen RR, Colman H, Flanders AE, Giannini C, Grifford M, Iavarone A, Jain R, Joseph I, Kim J, Kasaian K, Mikkelsen T, Murray BA, O’Neill BP, Pachter L, Parsons DW, Sougnez C, Sulman EP, Vandenberg SR, Van Meir EG, von Deimling A, Zhang H, Crain D, Lau K, Mallery D, Morris S, Paulauskis J, Penny R, Shelton T, Sherman M, Yena P, Black A, Bowen J, Dicostanzo K, Gastier-Foster J, Leraas KM, Lichtenberg TM, Pierson CR, Ramirez NC, Taylor C, Weaver S, Wise L, Zmuda E, Davidsen T, Demchok JA, Eley G, Ferguson ML, Hutter CM, Mills Shaw KR, Ozenberger BA, Sheth M, Sofia HJ, Tarnuzzer R, Wang Z, Yang L, Zenklusen JC, Ayala B, Baboud J, Chudamani S, Jensen MA, Liu J, Pihl T, Raman R, Wan Y, Wu Y, Ally A, Auman JT, Balasundaram M, Balu S, Baylin SB, Beroukhim R, Bootwalla MS, Bowlby R, Bristow CA, Brooks D, Butterfield Y, Carlsen R, Carter S, Chin L, Chu A, Chuah E, Cibulskis K, Clarke A, Coetzee SG, Dhalla N, Fennell T, Fisher S, Gabriel S, Getz G, Gibbs R, Guin R, Hadjipanayis A, Hayes DN, Hinoue T, Hoadley K, Holt RA, Hoyle AP, Jefferys SR, Jones S, Jones CD, Kucherlapati R, Lai PH, Lander E, Lee S, Lichtenstein L, Ma Y, Maglinte DT, Mahadeshwar HS, Marra MA, Mayo M, Meng S, Meyerson ML, Mieczkowski PA, Moore RA, Mose LE, Mungall AJ, Pantazi A, Parfenov M, Park PJ, Parker JS, Perou CM, Protopopov A, Ren X, Roach J, Sabedot TS, Schein J, Schumacher SE, Seidman JG, Seth S, Shen H, Simons JV, Sipahimalani P, Soloway MG, Song X, Sun H, Tabak B, Tam A, Tan D, Tang J, Thiessen N, Triche T Jr, Van Den Berg DJ, Veluvolu U, Waring S, Weisenberger DJ, Wilkerson MD, Wong T, Wu J, Xi L, Xu AW, Yang L, Zack TI, Zhang J, Aksoy BA, Arachchi H, Benz C, Bernard B, Carlin D, Cho J, DiCara D, Frazer S, Fuller GN, Gao J, Gehlenborg N, Haussler D, Heiman DI, Iype L, Jacobsen A, Ju Z, Katzman S, Kim H, Knijnenburg T, Kreisberg RB, Lawrence MS, Lee W, Leinonen K, Lin P, Ling S, Liu W, Liu Y, Liu Y, Lu Y, Mills G, Ng S, Noble MS, Paull E, Rao A, Reynolds S, Saksena G, Sanborn Z, Sander C, Schultz N, Senbabaoglu Y, Shen R, Shmulevich I, Sinha R, Stuart J, Sumer SO, Sun Y, Tasman N, Taylor BS, Voet D, Weinhold N, Weinstein JN, Yang D, Yoshihara K, Zheng S, Zhang W, Zou L, Abel T, Sadeghi S, Cohen ML, Eschbacher J, Hattab EM, Raghunathan A, Schniederjan MJ, Aziz D, Barnett G, Barrett W, Bigner DD, Boice L, Brewer C, Calatozzolo C, Campos B, Carlotti CG Jr, Chan TA, Cuppini L, Curley E, Cuzzubbo S, Devine K, DiMeco F, Duell R, Elder JB, Fehrenbach A, Finocchiaro G, Friedman W, Fulop J, Gardner J, Hermes B, Herold-Mende C, Jungk C, Kendler A, Lehman NL, Lipp E, Liu O, Mandt R, McGraw M, Mclendon R, McPherson C, Neder L, Nguyen P, Noss A, Nunziata R, Ostrom QT, Palmer C, Perin A, Pollo B, Potapov A, Potapova O, Rathmell WK, Rotin D, Scarpace L, Schilero C, Senecal K, Shimmel K, Shurkhay V, Sifri S, Singh R, Sloan AE, Smolenski K, Staugaitis SM, Steele R, Thorne L, Tirapelli DP, Unterberg A, Vallurupalli M, Wang Y, Warnick R, Williams F, Wolinsky Y, Bell S, Rosenberg M, Stewart C, Huang F, Grimsby JL, Radenbaugh AJ, Zhang J (2015) Comprehensive, integrative genomic analysis of diffuse lower-grade gliomas. N Engl J Med 372(26): 2481–2498. doi: 10.1056/NEJMoa1402121

9. Carriere PP, Haisraely O, Aaroe A, Ahmed K, Lewis R, Colson-Fearon D, Bolden M, Swanson TA, Beckham TH, Wang C, De B, Perni S, Tom MC, Li J, McGovern S, McAleer MF, Ghia A, Jiang W, Chung C, Grosshans D, Ballester L, Esquenazi Y, Kamiya-Matsuoka C, Yeboa DN (2026) Long-term outcomes in IDH-wildtype (IDH-wt) gliomas with historical WHO grade 2 and 3 histology. J Neurooncol 179(1): 12. doi: 10.1007/s11060-026-05712-2

10. Chang SM, Cahill DP, Aldape KD, Mehta MP (2016) Treatment of adult lower-grade glioma in the era of genomic medicine. Am Soc Clin Oncol Educ Book 35: 75–81. doi: 10.1200/EDBK_158869

11. Drexler R, Drinnenberg A, Gavish A, Yalçin B, Shamardani K, Rogers AE, Mancusi R, Trivedi V, Taylor KR, Kim YS, Woo PJ, Soni N, Su M, Ravel A, Tatlock E, Midler A, Wu SH, Ramakrishnan C, Chen R, Ayala-Sarmiento AE, Fernandez Pacheco DR, Siverts L, Daigle TL, Tasic B, Zeng H, Breunig JJ, Deisseroth K, Monje M (2025a) Cholinergic neuronal activity promotes diffuse midline glioma growth through muscarinic signaling. Cell 188(17): 4640–4657.e30. doi: 10.1016/j.cell.2025.05.031

12. Drexler R, Yalçın B, Mancusi R, Rogers A, Shamardani K, Woo PJ, Ravel A, Wu S, Yabo YA, Steger L, de Biagi-Junior CAO, Cascio CL, Malenka R, Heifets BD, Filbin MG, Heiland DH, Deisseroth K, Monje M (2025b) Serotonergic neuron-glioma interactions drive high-grade glioma pathophysiology. bioRxiv [Preprint] Dec 12:2025.12.10.693579. doi: 10.64898/2025.12.10.693579

13. Fu JQ, Chen Z, Hu YJ, Fan ZH, Guo ZX, Liang JY, Ryu BM, Ren JL, Shi XJ, Li J, Jia S, Wang J, Ke XS, Ma X, Tan X, Zhang T, Chen XZ, Zhang C (2019). A single factor induces neuronal differentiation to suppress glioma cell growth. CNS Neurosci Ther 25(4): 486–495. doi: 10.1111/cns.13066

14. Gaia F, Dal-Pizzol HR, Malafaia O, Roesler R, Isolan GR (2026) DLG2-DLG4 expression is associated with improved survival and a synaptic gene signature in lower-grade glioma. Cancers (Basel) 18(10): 1646. doi: 10.3390/cancers18101646

15. Grubor M, Zivkovic M, Sagud M, Nikolac Perkovic M, Mihaljevic-Peles A, Pivac N, Muck-Seler D, Svob Strac D (2020) HTR1A, HTR1B, HTR2A, HTR2C and HTR6 gene polymorphisms and extrapyramidal side effects in haloperidol-treated patients with schizophrenia. Int J Mol Sci 21(7): 2345. doi: 10.3390/ijms21072345

16. Gue R, Lakhani DA (2024) The 2021 World Health Organization Central Nervous System Tumor Classification: The spectrum of diffuse gliomas. Biomedicines 12(6): 1349. doi: 10.3390/biomedicines12061349

17. Gwynne WD, Shakeel MS, Girgis-Gabardo A, Hassell JA (2021) The role of serotonin in breast cancer stem cells. Molecules 26(11): 3171. doi: 10.3390/molecules26113171

18. Holmes C, Arranz M, Collier D, Powell J, Lovestone S (2003) Depression in Alzheimer’s disease: the effect of serotonin receptor gene variation. Am J Med Genet B Neuropsychiatr Genet 119B(1): 40-43. doi: 10.1002/ajmg.b.10068

19. Holmes C, Arranz MJ, Powell JF, Collier DA, Lovestone S (1998) 5-HT2A and 5-HT2C receptor polymorphisms and psychopathology in late onset Alzheimer’s disease. Hum Mol Genet 7(9): 1507–1509. doi: 10.1093/hmg/7.9.1507

20. Hoyer D, Martin G. 5-HT receptor classification and nomenclature: towards a harmonization with the human genome. Neuropharmacology. 1997 Apr-May;36(4-5):419-28. doi: 10.1016/s0028-3908(97)00036-1

21. Huang YY, Kandel ER (2007) 5-Hydroxytryptamine induces a protein kinase A/mitogen-activated protein kinase-mediated and macromolecular synthesis-dependent late phase of long-term potentiation in the amygdala. J Neurosci 27(12): 3111–3119. doi: 10.1523/JNEUROSCI.3908-06.2007

22. Illi A, Setälä-Soikkeli E, Viikki M, Poutanen O, Huhtala H, Mononen N, Lehtimäki T, Leinonen E, Kampman O (2009) 5-HTR1A, 5-HTR2A, 5-HTR6, TPH1 and TPH2 polymorphisms and major depression. Neuroreport 20(12): 1125–1128. doi: 10.1097/WNR.0b013e32832eb708

23. Ji Y, Ju CW, Chen L, Shen K, Su R, Li A, Liu X, Liu B, Zhang X, Lyu R, Xia P, Li H, Pan Y, Liu Y, Tse MH, Xue Y, Qian H, Jing N, Zhu HH, Wang L, Zhang LS, Jiang SH, Zhang W, Dong L, Yan Z, Pan J, Zhu Y, Wei J, Wang Q, Xue W (2026) Serotonin modulates lineage plasticity in neuroendocrine prostate cancer via epigenetic reprogramming. Cancer Discov 16(4): 760–780. doi: 10.1158/2159-8290.CD-25-0974

24. Johnson KC, Spitzer A, Varn FS, Nomura M, Garofano L, Chowdhury T, Lipsa A, Zhang L, Fernández EC, Barak T, Gulhan Ercan-Sencicek A, Peksen AB, Anderson KJ, Tesileanu CMS, Amin SB, Kocakavuk E, Zhao D, D’Angelo F, Migliozzi S, Bussema L, Gritsch S, Moon HE, Paek SH, Bielle F, Laurenge A, Di Stefano AL, Mathon B, Picca A, Sanson M, Hau AC, Hertel F, Grzyb K, Zhao Z, Wang Q, Jiang T, Miller JJ, Wakimoto H, Cahill DP, Moliterno J, Günel M, Hermes B, Sanai N, Golebiewska A, Niclou SP, Huse J, Alfred Yung WK, Lasorella A, Suvà ML, Iavarone A, Tirosh I, Verhaak RGW (2026) Acquired genetic and cell-state changes in IDH-mutant glioma progression. Nature 655(8124): 1048–1059. doi: 10.1038/s41586-026-10612-6

25. Julius D, MacDermott AB, Axel R, Jessell TM (1988). Molecular characterization of a functional cDNA encoding the serotonin 1c receptor. Science 241(4865): 558–564. doi: 10.1126/science.3399891

26. Kihlstedt CJ, Dénes A, Mansouri A, Mikolajewicz N, Skoglund T, Köster L, Corell A, Carén H, Ferreyra Vega S, Olsson Bontell T, Thorsell A, Jakola AS (2025) Proteomics in IDH-mutant diffuse lower-grade glioma: a scoping review. Neurooncol Adv 8(1): vdaf258. doi: 10.1093/noajnl/vdaf258

27. Klempin F, Babu H, De Pietri Tonelli D, Alarcon E, Fabel K, Kempermann G (2010) Oppositional effects of serotonin receptors 5-HT1a, 2, and 2c in the regulation of adult hippocampal neurogenesis. Front Mol Neurosci 3:14. doi: 10.3389/fnmol.2010.00014

28. Kobilka BK, Frielle T, Collins S, Yang-Feng T, Kobilka TS, Francke U, Lefkowitz RJ, Caron MG (1987) An intronless gene encoding a potential member of the family of receptors coupled to guanine nucleotide regulatory proteins. Nature 329(6134): 75–79. doi: 10.1038/329075a0

29. Kolan SS, Lidström T, Mediavilla T, Dernstedt A, Degerman S, Hultdin M, Björk K, Marcellino D, Forsell MNE (2019) Growth-inhibition of cell lines derived from B cell lymphomas through antagonism of serotonin receptor signaling. Sci Rep 9(1): 4276. doi: 10.1038/s41598-019-40825-x

30. Li J, Lu J, Zheng C, Huang X, Li H, Mai Q, Chen S, Zhou Z, Zhu J, Yu T, Xu M, Tan H, Zhang CM, Gao Q, Liu J, Pan C (2026) Serotonin-licensed macrophages potentiate chemoresistance via inositol metabolic crosstalk in ovarian cancer. Cell Metab 2026 38(2): 331-349.e10. doi: 10.1016/j.cmet.2025.11.011

31. Louis DN, Perry A, Wesseling P, Brat DJ, Cree IA, Figarella-Branger D, Hawkins C, Ng HK, Pfister SM, Reifenberger G, Soffietti R, von Deimling A, Ellison DW (2021) The 2021 WHO Classification of Tumors of the Central Nervous System: A summary. Neuro Oncol 23(8): 1231–1251. doi: 10.1093/neuonc/noab106.

32. Messina D, Annesi G, Serra P, Nicoletti G, Pasqua A, Annesi F, Tomaino C, Cirò-Candiano IC, Carrideo S, Caracciolo M, Spadafora P, Zappia M, Savettieri G, Quattrone A (2002) Association of the 5-HT6 receptor gene polymorphism C267T with Parkinson’s disease. Neurology 58(5): 828–829. doi: 10.1212/wnl.58.5.828

33. Monje M (2025) The neuroscience of brain cancers. Neuron 113(17): 2734–2739. doi: 10.1016/j.neuron.2025.07.012

34. Monsma FJ Jr, Shen Y, Ward RP, Hamblin MW, Sibley DR (1993) Cloning and expression of a novel serotonin receptor with high affinity for tricyclic psychotropic drugs. Mol Pharmacol 43(3):320–327.

35. Ogelman R, Gomez Wulschner LE, Hoelscher VM, Hwang IW, Chang VN, Oh WC (2024) Serotonin modulates excitatory synapse maturation in the developing prefrontal cortex. Nat Commun 15(1): 1368. doi: 10.1038/s41467-024-45734-w

36. Pritchard AL, Harris J, Pritchard CW, Coates J, Haque S, Holder R, Bentham P, Lendon CL (2008) Role of 5HT 2A and 5HT 2C polymorphisms in behavioural and psychological symptoms of Alzheimer’s disease. Neurobiol Aging 29(3): 341–347. doi: 10.1016/j.neurobiolaging.2006.10.011

37. Pritchett DB, Bach AW, Wozny M, Taleb O, Dal Toso R, Shih JC, Seeburg PH (1988) Structure and functional expression of cloned rat serotonin 5HT-2 receptor. EMBO J 7(13): 4135–4140. doi: 10.1002/j.1460-2075.1988.tb03308.x

38. Rodrigues B, Dalmolin M, Dal-Pizzol HR, Malafaia O, Fernandes MAC, Coelho KMdPA, Roesler R, Isolan GR (2026) AMPAR subunit gene expression marks a synaptic transcriptional state in lower-grade glioma. Brain Sciences 16(8): 773. 10.3390/brainsci16080773

39. Saeb L, Abtahi S, Masoudi R, Javadpour A (2026) Association of polymorphisms in genes involved in serotonergic signaling with the risk of developing Alzheimer’s disease. Curr Alzheimer Res doi: 10.2174/0115672050459074260331053012

40. Shen J, Wang Q, Lu F, Xu H, Wang P, Feng Y (2023) Prognostic and immunomodulatory roles of schizophrenia-associated genes HTR2A, COMT, and PRODH in pan-cancer analysis and glioma survival prediction model. Front Immunol 14: 1201252. doi: 10.3389/fimmu.2023.1201252

41. Tan YF, Yeh CY, Hsu SY, Lu CH, Tsai CH, Chiang PC, Weng HJ, Tsai TF, Lee YL (2025) Serotonin 2A receptor attenuates psoriatic inflammation by suppressing IL-23 secretion in monocyte-derived Langerhans cells. Nat Commun 16(1): 8544. doi: 10.1038/s41467-025-63971-5

42. Taylor KR, Barron T, Hui A, Spitzer A, Yalçin B, Ivec AE, Geraghty AC, Hartmann GG, Arzt M, Gillespie SM, Kim YS, Maleki Jahan S, Zhang H, Shamardani K, Su M, Ni L, Du PP, Woo PJ, Silva-Torres A, Venkatesh HS, Mancusi R, Ponnuswami A, Mulinyawe S, Keough MB, Chau I, Aziz-Bose R, Tirosh I, Suvà ML, Monje M (2023) Glioma synapses recruit mechanisms of adaptive plasticity. Nature 623(7986): 366–374. doi: 10.1038/s41586-023-06678-1

43. Thome J, Retz W, Baader M, Pesold B, Hu M, Cowen M, Durany N, Adler G, Henn FA, Rösler M (2001) Association analysis of HTR6 and HTR2A polymorphisms in sporadic Alzheimer’s disease. J Neural Transm (Vienna) 108(10): 1175–1180. doi: 10.1007/s007020170007

44. Tirosh I, Venteicher AS, Hebert C, Escalante LE, Patel AP, Yizhak K, Fisher JM, Rodman C, Mount C, Filbin MG, Neftel C, Desai N, Nyman J, Izar B, Luo CC, Francis JM, Patel AA, Onozato ML, Riggi N, Livak KJ, Gennert D, Satija R, Nahed BV, Curry WT, Martuza RL, Mylvaganam R, Iafrate AJ, Frosch MP, Golub TR, Rivera MN, Getz G, Rozenblatt-Rosen O, Cahill DP, Monje M, Bernstein BE, Louis DN, Regev A, Suvà ML (2016) Single-cell RNA-seq supports a developmental hierarchy in human oligodendroglioma. Nature 539(7628): 309–313. doi: 10.1038/nature20123

45. Udoh UG, Bruno JR, Osborn PO, Pratt KG (2024) Serotonin strengthens a developing glutamatergic synapse through a PI3K-dependent mechanism. J Neurosci 44(6): e1260232023. doi: 10.1523/JNEUROSCI.1260-23.2023

46. Upreti C, Konstantinov E, Kassabov SR, Bailey CH, Kandel ER (2019) Serotonin induces structural plasticity of both extrinsic modulating and intrinsic mediating circuits in vitro in Aplysia californica. Cell Rep 28(11): 2955–2965.e3. doi: 10.1016/j.celrep.2019.08.016

47. Venkataramani V, Tanev DI, Strahle C, Studier-Fischer A, Fankhauser L, Kessler T, Körber C, Kardorff M, Ratliff M, Xie R, Horstmann H, Messer M, Paik SP, Knabbe J, Sahm F, Kurz FT, Acikgöz AA, Herrmannsdörfer F, Agarwal A, Bergles DE, Chalmers A, Miletic H, Turcan S, Mawrin C, Hänggi D, Liu HK, Wick W, Winkler F, Kuner T (2019) Glutamatergic synaptic input to glioma cells drives brain tumour progression. Nature 573(7775): 532–538. doi: 10.1038/s41586-019-1564-x

48. Venkatesh HS, Johung TB, Caretti V, Noll A, Tang Y, Nagaraja S, Gibson EM, Mount CW, Polepalli J, Mitra SS, Woo PJ, Malenka RC, Vogel H, Bredel M, Mallick P, Monje M (2015) Neuronal activity promotes glioma growth through neuroligin-3 secretion. Cell 161(4): 803–816. doi: 10.1016/j.cell.2015.04.012

49. Venkatesh HS, Morishita W, Geraghty AC, Silverbush D, Gillespie SM, Arzt M, Tam LT, Espenel C, Ponnuswami A, Ni L, Woo PJ, Taylor KR, Agarwal A, Regev A, Brang D, Vogel H, Hervey-Jumper S, Bergles DE, Suvà ML, Malenka RC, Monje M (2019) Electrical and synaptic integration of glioma into neural circuits. Nature 573(7775): 539–545. doi: 10.1038/s41586-019-1563-y

50. Venteicher AS, Tirosh I, Hebert C, Yizhak K, Neftel C, Filbin MG, Hovestadt V, Escalante LE, Shaw ML, Rodman C, Gillespie SM, Dionne D, Luo CC, Ravichandran H, Mylvaganam R, Mount C, Onozato ML, Nahed BV, Wakimoto H, Curry WT, Iafrate AJ, Rivera MN, Frosch MP, Golub TR, Brastianos PK, Getz G, Patel AP, Monje M, Cahill DP, Rozenblatt-Rosen O, Louis DN, Bernstein BE, Regev A, Suvà ML (2017) Decoupling genetics, lineages, and microenvironment in IDH-mutant gliomas by single-cell RNA-seq. Science 355(6332): eaai8478. doi: 10.1126/science.aai8478

51. Wouters MM, Roeder JL, Tharayil VS, Stanich JE, Strege PR, Lei S, Bardsley MR, Ordog T, Gibbons SJ, Farrugia G. Protein kinase C{gamma} mediates regulation of proliferation by the serotonin 5-hydroxytryptamine receptor 2B. J Biol Chem. 2009 Aug 7;284(32):21177–84. doi: 10.1074/jbc.M109.015859

52. Xing L, Kalebic N, Namba T, Vaid S, Wimberger P, Huttner WB (2020) Serotonin receptor 2A activation promotes evolutionarily relevant basal progenitor proliferation in the developing neocortex. Neuron 108(6): 1113–1129.e6. doi: 10.1016/j.neuron.2020.09.034

53. Yi Y, Che W, Xu P, Mao C, Li Z, Wang Q, Lyu J, Wang X (2024) Conversion of glioma cells into neuron-like cells by small molecules. iScience 27(11): 111091. doi: 10.1016/j.isci.2024.111091

54. Yoshihara K, Shahmoradgoli M, Martínez E, Vegesna R, Kim H, Torres-Garcia W, Treviño V, Shen H, Laird PW, Levine DA, Carter SL, Getz G, Stemke-Hale K, Mills GB, Verhaak RG (2013) Inferring tumour purity and stromal and immune cell admixture from expression data. Nat Commun 4: 2612. doi: 10.1038/ncomms3612

55. Zamani A, Qu Z (2012) Serotonin activates angiogenic phosphorylation signaling in human endothelial cells. FEBS Lett 586(16): 2360–2365. doi: 10.1016/j.febslet.2012.05.047

56. Zhan D, Wang X, Zheng Y, Wang S, Yang B, Pan B, Wang N, Wang Z (2023) Integrative dissection of 5-hydroxytryptamine receptors-related signature in the prognosis and immune microenvironment of breast cancer. Front Oncol 13: 1147189. doi: 10.3389/fonc.2023.1147189

57. Zhang W, Li L, Li J, Yu H, Zheng F, Yan B, Cai W, Chen Y, Yin L, Tang D, Xu Y, Dai Y (2022) Systematic analysis of neurotransmitter receptors in human breast cancer reveals a strong association with outcome and uncovers HTR6 as a survival-associated gene potentially regulating the immune microenvironment. Front Immunol 13: 756928. doi: 10.3389/fimmu.2022.756928

58. Zhao Z, Zhang KN, Wang Q, Li G, Zeng F, Zhang Y, Wu F, Chai R, Wang Z, Zhang C, Zhang W, Bao Z, Jiang T (2021) Chinese Glioma Genome Atlas (CGGA): A comprehensive resource with functional genomic data from Chinese glioma patients. Genomics Proteomics Bioinformatics 19(1): 1–12. doi: 10.1016/j.gpb.2020.10.005

