## Supplementary Information for "A Subset of G Protein-Coupled Serotonin Receptor Genes is Linked to a Neuronal Gene Expression Signature and Clinically Favorable Biology in IDH-Mutant Gliomas"

**Supplementary Table S1.** Statistics for comparisons in expression of *HTR1A*, *HTR2A*, *HTR2C*, and *HTR6* in TCGA LGG tumors according to the mutational status of IDH1, IDH2, and glioma driver genes.

| | **Diff** | **Lower** | **Upper** | **Adjusted *P*-value** | **Significance** |
|---|---|---|---|---|---|
| ***HTR1A*** | | | | | |
| **IDH1 status** | | | | | |
| Mutated-Wild-type | 1.24 | 0.63 | 1.85 | < 0.001 | *** |
| **IDH2 status** | | | | | |
| Mutated-Wild-type | 2.15 | 0.85 | 3.45 | < 0.01 | ** |
| ***TP53* status** | | | | | |
| Mutated-Wild-type | -0.61 | -1.13 | -0.08 | 0.02 | * |
| ***PTEN* status** | | | | | |
| Mutated-Wild-type | -2.84 | -4.07 | -1.61 | < 0.001 | *** |
| ***EGFR* status** | | | | | |
| Mutated-Wild-type | -1.97 | -3.10 | -0.84 | < 0.001 | *** |
| ***HTR2A*** | | | | | |
| **IDH1 status** | | | | | |
| Mutated-Wild-type | 0.84 | 0.34 | 1.34 | < 0.01 | ** |
| **IDH2 status** | | | | | |
| Mutated-Wild-type | 1.28 | 0.21 | 2.35 | 0.02 | * |
| ***TP53* status** | | | | | |
| Mutated-Wild-type | -0.67 | -1.10 | -0.24 | < 0.01 | ** |
| ***PTEN* status** | | | | | |
| Mutated-Wild-type | -1.72 | -2.74 | -0.69 | < 0.01 | ** |
| ***EGFR* status** | | | | | |
| Mutated-Wild-type | -1.23 | -2.17 | -0.30 | 0.01 | * |
| ***HTR2C*** | | | | | |
| **IDH1 status** | | | | | |
| Mutated-Wild-type | 0.27 | -0.30 | 0.85 | 0.35 | NS |
| **IDH2 status** | | | | | |
| Mutated-Wild-type | 1.51 | 0.30 | 2.73 | 0.02 | * |
| ***TP53* status** | | | | | |
| Mutated-Wild-type | -0.39 | -0.88 | 0.10 | 0.12 | NS |
| ***PTEN* status** | | | | | |
| Mutated-Wild-type | -2.14 | -3.30 | -0.99 | < 0.001 | *** |

| | | | | | |
|---|---|---|---|---|---|
| ***EGFR* status** | | | | | |
| Mutated-Wild-type | -1.49 | -2.55 | -0.43 | < 0.01 | ** |
| | | ***HTR6*** | | | |
| **IDH1 status** | | | | | |
| Mutated-Wild-type | 0.10 | -0.39 | 0.58 | 0.70 | NS |
| **IDH2 status** | | | | | |
| Mutated-Wild-type | 2.14 | 1.14 | 3.15 | < 0.001 | *** |
| ***TP53* status** | | | | | |
| Mutated-Wild-type | -1.18 | -1.57 | -0.79 | < 0.001 | *** |
| ***PTEN* status** | | | | | |
| Mutated-Wild-type | -1.34 | -2.32 | -0.35 | < 0.01 | ** |
| ***EGFR* status** | | | | | |
| Mutated-Wild-type | -0.70 | -1.60 | 0.20 | 0.13 | NS |

Diff, estimated difference between the mean expression values of the groups being compared. Lower, lower boundary of the 95% confidence interval for that mean difference. Upper, upper boundary of the 95% confidence interval for that mean difference. NS, non-significant.

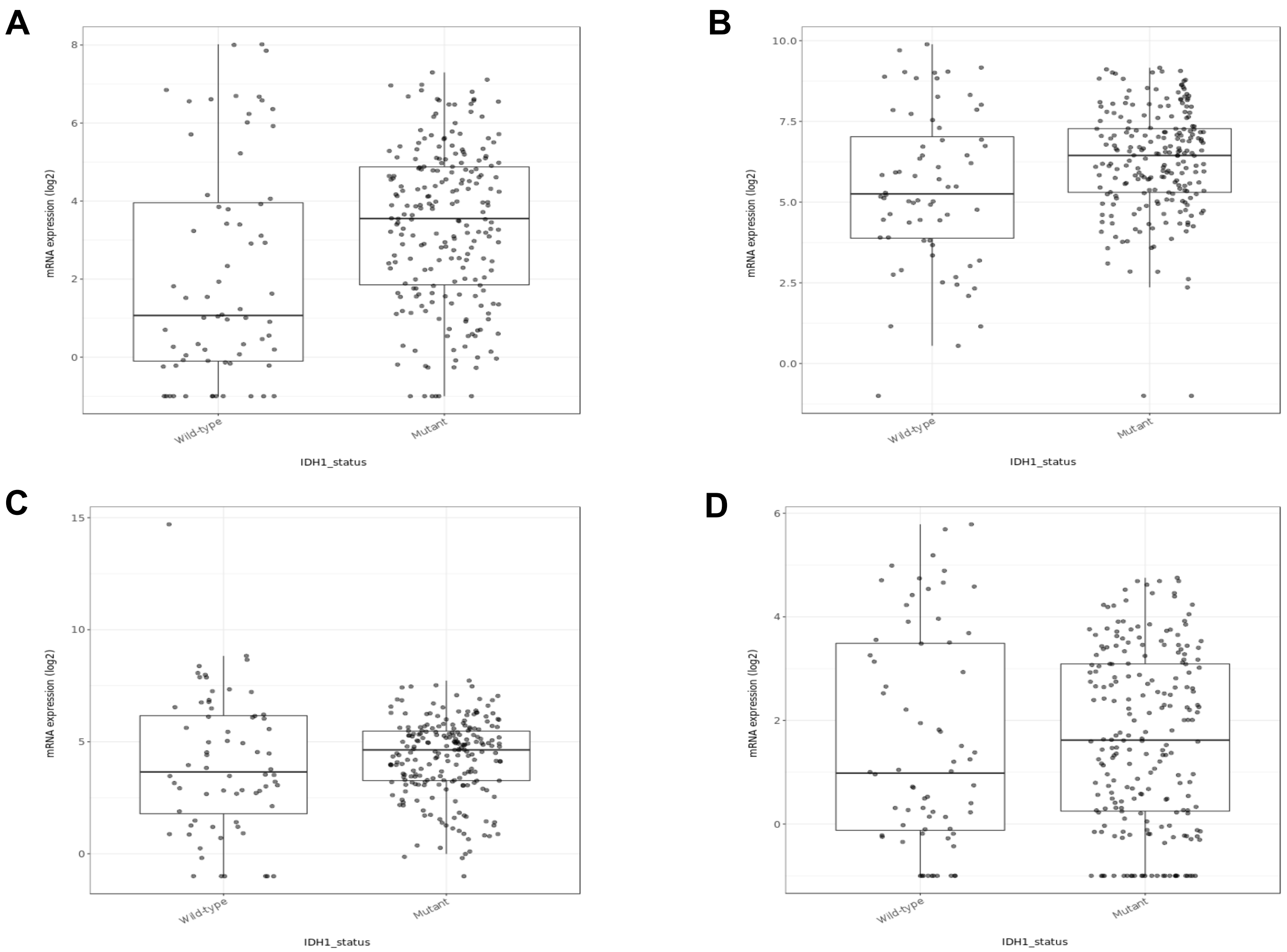

**Supplementary Fig. S1.** Expression of 5-HT receptor genes according to IDH1 status in TCGA-LGG tumors. Expression levels of *HTR1A* (**A**), *HTR2A* (**B**), *HTR2C* (**C**), and *HTR6* (**D**) in IDH1-wild-type and IDH1-mutated tumors; *n* = 286 tumors for which gene expression was matched with mutation status in Gliovis (https://gliovis.bioinfo.cnio.es/). Statistical comparison results are summarized in Supplementary Table S1.

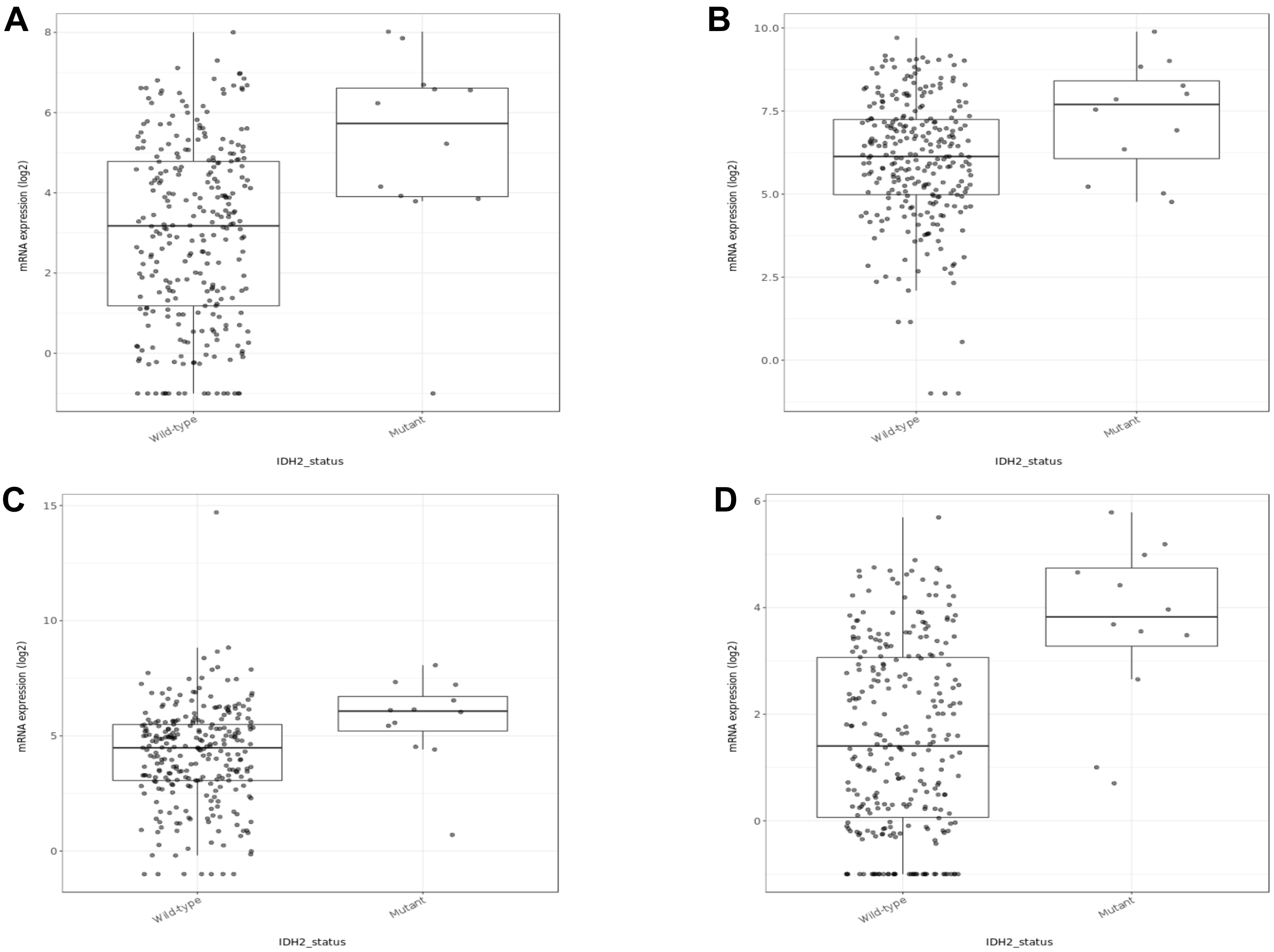

**Supplementary Fig. S2.** Expression of 5-HT receptor genes according to IDH2 status in TCGA-LGG tumors. Expression levels of *HTR1A* (**A**), *HTR2A* (**B**), *HTR2C* (**C**), and *HTR6* (**D**) in IDH2-wild-type and IDH2-mutated tumors; $n$ = 286 tumors for which gene expression was matched with mutation status in Gliovis (https://gliovis.bioinfo.cnio.es/). Statistical comparison results are summarized in Supplementary Table S1.

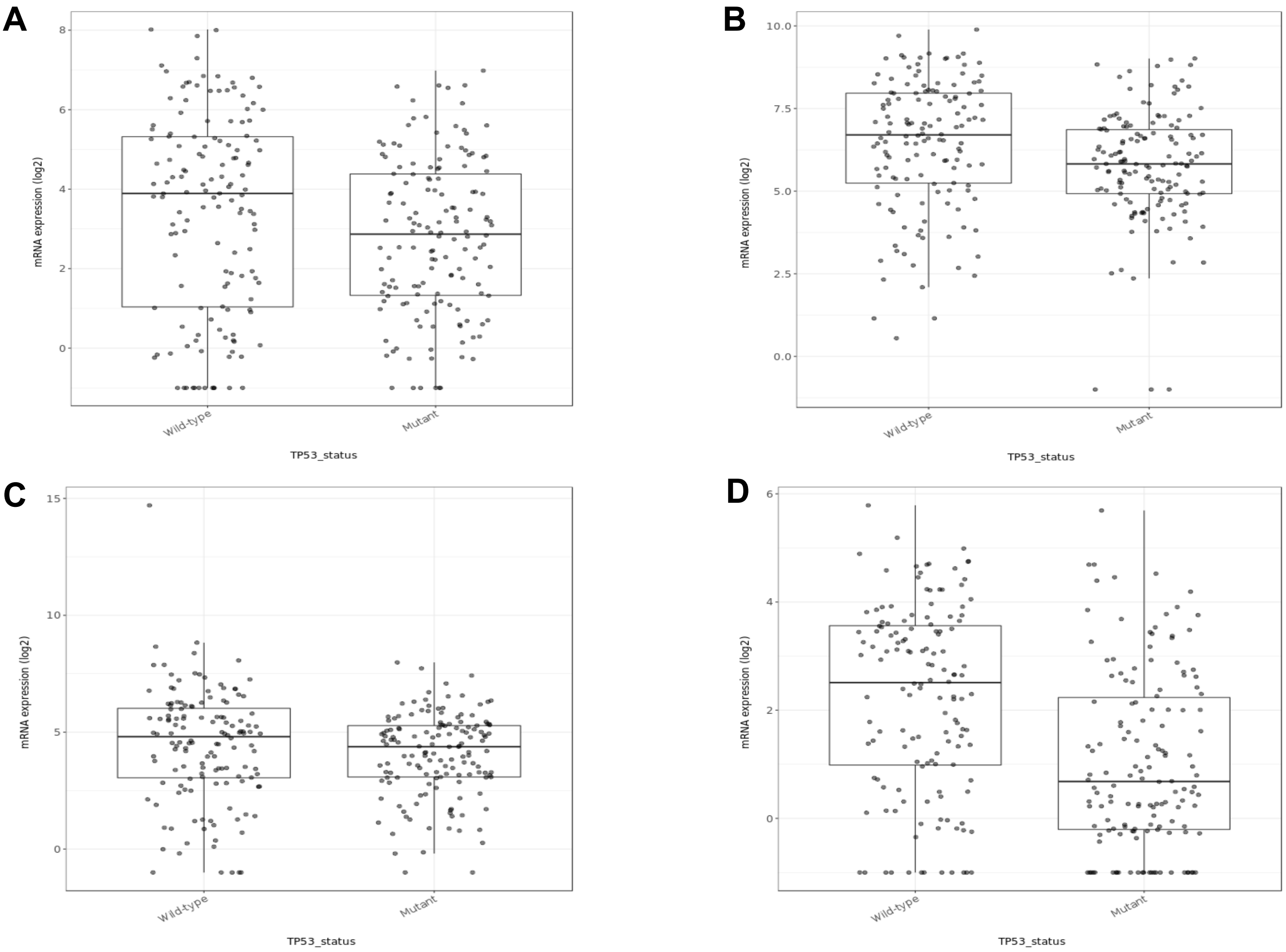

**Supplementary Fig. S3.** Expression of 5-HT receptor genes according to *TP53* status in TCGA-LGG tumors. Expression levels of *HTR1A* (**A**), *HTR2A* (**B**), *HTR2C* (**C**), and *HTR6* (**D**) in *TP53*-wild-type and *TP53*-mutated tumors; *n* = 286 tumors for which gene expression was matched with mutation status in Gliovis (https://gliovis.bioinfo.cnio.es/). Statistical comparison results are summarized in Supplementary Table S1.

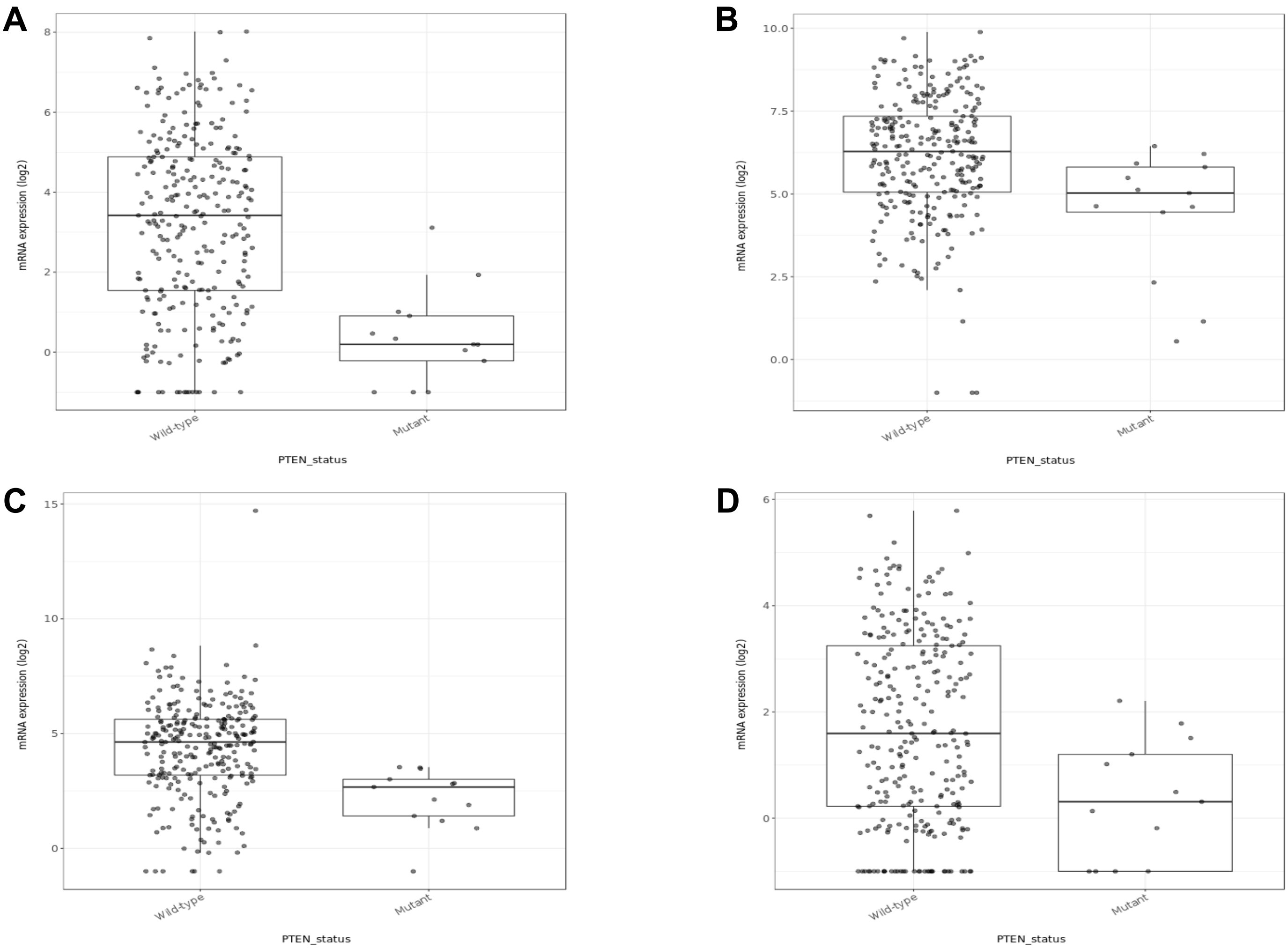

**Supplementary Fig. S4.** Expression of 5-HT receptor genes according to *PTEN* status in TCGA-LGG tumors. Expression levels of *HTR1A* (**A**), *HTR2A* (**B**), *HTR2C* (**C**), and *HTR6* (**D**) in *PTEN*-wild-type and *PTEN*-mutated tumors; $n$ = 286 tumors for which gene expression was matched with mutation status in Gliovis (https://gliovis.bioinfo.cnio.es/). Statistical comparison results are summarized in Supplementary Table S1.

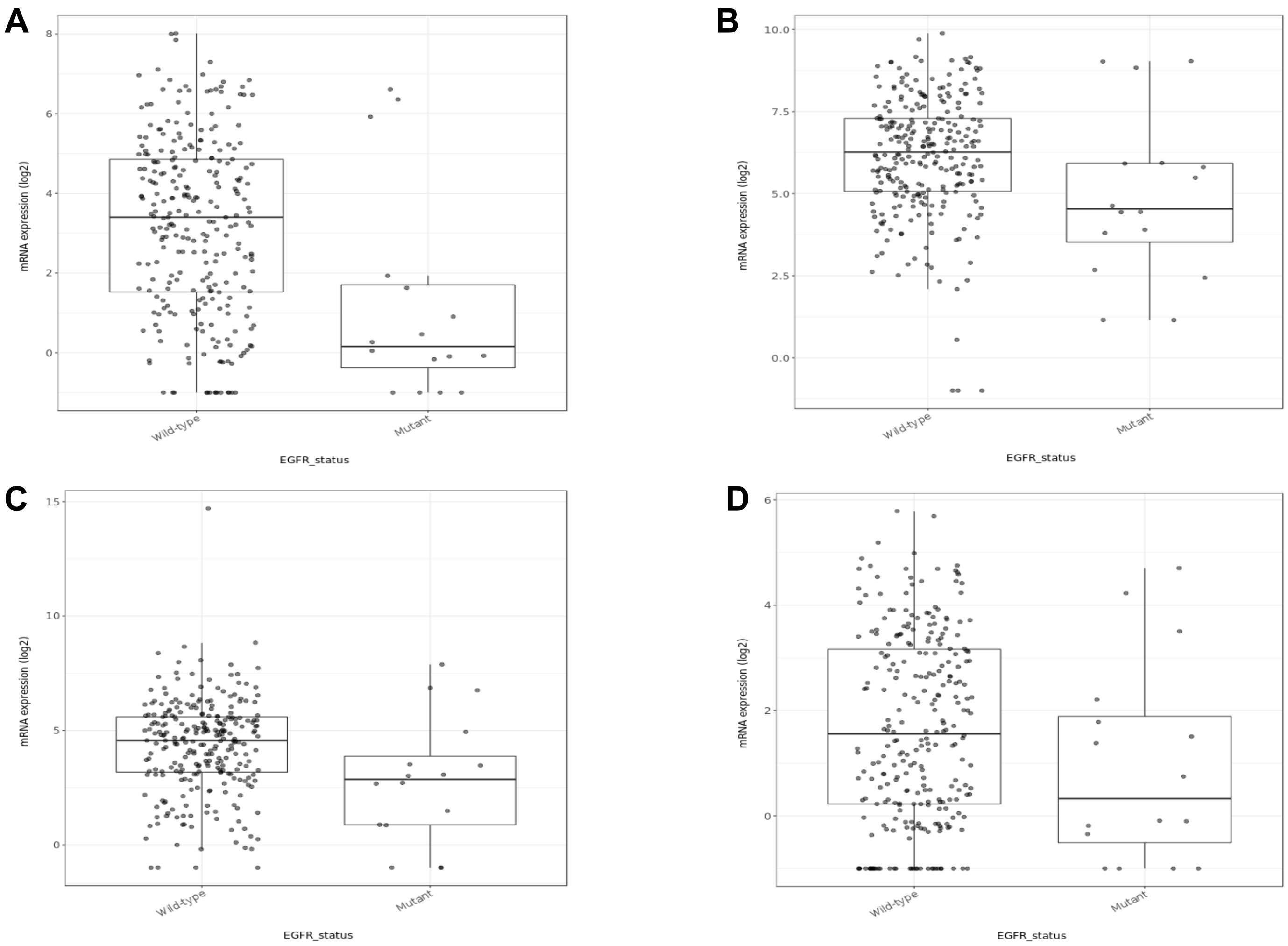

**Supplementary Fig. S5.** Expression of 5-HT receptor genes according to *EGFR* status in TCGA-LGG tumors. Expression levels of *HTR1A* (**A**), *HTR2A* (**B**), *HTR2C* (**C**), and *HTR6* (**D**) in *EGFR*-wild-type and *EGFR*-mutated tumors; *n* = 286 tumors for which gene expression was matched with mutation status in Gliovis (https://gliovis.bioinfo.cnio.es/). Statistical comparison results are summarized in Supplementary Table S1.
